# Nuclear transport under stress phenocopies transport defects in models of C9Orf72 ALS

**DOI:** 10.1101/2022.04.13.488135

**Authors:** Marije F.W. Semmelink, Hamidreza Jafarinia, Justina C Wolters, Teodora Gheorghe, Sara N. Mouton, Anton Steen, Patrick R. Onck, Liesbeth M. Veenhoff

## Abstract

The nucleus is the hallmark of eukaryotic life and transport to and from the nucleus occurs through the nuclear pore complex (NPC). There is a multitude of data connecting the nuclear transport machinery – i.e. the NPCs and associated nuclear transport factors - to neurodegenerative diseases, but the mechanisms are not well understood. Using *Saccharomyces cerevisiae*, we systematically studied how the expression of polyPR and polyGA related to C9Orf72 amyotrophic lateral sclerosis impacts the nuclear transport machinery. We measured the abundance and localization of NPC components and transport factors, and assessed the kinetics of import and export by four transport receptors. PolyPR and polyGA cause distinct, and transport receptor dependent effects. We compared the specific changes in transport to those obtained when cells were exposed to different stress situations or mutations. This comparison showed similar patterns of transport defects in cells lacking specific NTRs and cells expressing polyPR. In contrast, polyGA expressing cells bear resemblance to stress conditions where energy maintenance is decreased. The similarity of the patterns of transport deficiencies suggests that polyPR has a direct effect on nuclear transport via NTRs, while polyGA impacts the energy state of the cell and subsequently changes transport.

## Introduction

### Nucleocytoplasmic transport

For all eukaryotes, DNA is safely stored in their nuclei, which allows for compartmentalization of transcription and translation, thereby contributing to the control of gene expression. The nuclear envelope separates the contents of the nucleus from the cytoplasm, and contains nuclear pore complexes (NPCs) which perform highly regulated transport as well as passive diffusion through their permeability barrier. The NPCs are large protein complexes build from nucleoporins (Nups), with roughly half of the mass of an NPC constituted by Nups containing phenylalanine-glycine repeats (FG-Nups)^1^. Additionally, a pool of Importin-β functions as a stable component of the NPC’s permeability barrier^2^. Nucleocytoplasmic transport (NCT) through the pores is performed by several nuclear transport receptors (NTRs) and together they bind a large and diverse group of cargoes. While general protein export is mediated only by the major exportin Crm1, different importins are jointly responsible for all protein import. Baker’s yeast contains 18 known NTRs, whereas in human cells 30 NTRs have been identified^3–7^. The Importin-β superfamily is the largest class of NTRs; they can either directly bind to their cargo, or via an adaptor protein, an Importin-α isoform^8^. Cargo recognition is achieved via either a nuclear localization signal (NLS) which binds to an importin, or a nuclear export signal (NES) which binds to an exportin. For the NTRs some, but not all, cargoes are known, and redundancies exist^9^. The directionality of the transport is maintained by a gradient of Ran, a member of the Ras family of small GTPases, which is bound to GTP in the nucleus and to GDP in the cytoplasm. Importins require RanGTP to dissociate from their cargo in the nucleus, and exportins require RanGTP to bind simultaneously with their cargo to allow binding^10–12^.

### Transport in neurodegeneration and under stress conditions

Nuclear transport of transcriptional regulators is a key step in responding to stress. For example, it was shown that transport by specific NTRs increases or decreases in response to heat shock^13–16^, glucose deprivation^17–20^, osmotic shock^21,22^, or oxidative stress^15,23–26^. Neurodegenerative diseases are also linked to alterations in NCT. This may be indirect, for example via the mentioned stress responses, or the connection may be more direct through interactions of the disease proteins with the NPC components or nuclear transport factors. E.g. mutant Huntingtin, co-aggregates Nups and RanGAP, reduces the abundance of nuclear RanGTP, and reduces protein import and export^27^. Alzheimer’s related tau protein also mislocalizes Nups and Ran, changes NPC permeability, and protein import and export^28^. The mutant form of TDP-43 (TAR DNA-binding protein-43) aggregates and is linked to ALS. Its aggregation triggers the mislocalization of Nups and NTRs, and interferes with protein import and RNA export^29^. Mutations in the NLS of the Fused in Sarcoma (FUS) protein reduce binding to its importin, and cause aggregation in cytoplasmic stress granules in FUS-related ALS^30^. Alteration of protein import and export was also seen in the presence of non-natural, synthetic β-sheet proteins, which suggests protein aggregation in general may be linked to reduced NCT^31^.

A specific type of ALS is caused by a repeat-expansion of G_4_C_2_ in *C9orf72* (C9ALS), which leads to disruption of the C9orf72 intron and thus a loss of function of the C9orf72 protein. In addition, the repeat RNA may cause a toxic gain of function, and, the extended G_4_C_2_ repeats leads to unconventional repeat-<u>a</u>ssociated <u>n</u>on-AUG (RAN) translation. This non-canonical initiation of translation allows elongation of a repeat sequence in the absence of an AUG starting codon, in multiple reading frames, and so generates multiple dipeptide repeat proteins (DPRs)^32^. For *C9orf72*, the sense RNA encodes glycine-alanine (GA), glycine-proline (GP), and glycine-arginine (GR) repeat proteins. The antisense transcript produces proline-arginine (PR), GP, and proline-alanine (PA)^33,34^ repeat proteins. Of these DPRs, polyPR is the most toxic species in most studied models^35–39^, followed by polyGR, while polyGA is toxic in about half the models, and polyPA is never toxic (reviewed in^40^).

The underlying cause for toxicity in C9ALS is still unknown, but interaction with NCT was shown in multiple studies (reviewed in^41–45^). The C9orf72 protein was found to interact with both Importin β1 and Ran-GTPase^46^. (G_4_C_2_)_30_ RNA was shown to directly bind to RanGAP^47^, and polyGA was reported to bind RanGAP1 in cytoplasmic inclusions^48^, and to reduce NCT^49^. The toxicity of (G_4_C_2_)_58_ was further linked to impaired RNA export, and moreover knockout or overexpression of many nuclear transport proteins was found to influence neurodegeneration^50^. Direct binding of PR to one of the NTRs, importin-β, resulted in decreased nuclear import of importin β, importin α/β, and transportin cargoes in permeabilized mouse neurons and HeLa cells^51^. In addition, nuclear envelopes incubated with PR_20_ peptides showed polyPR presence in the main channel of NPCs^52^. Furthermore, the distribution of protein components of NPCs were found to be altered in ALS patients and ALS mouse models^53^. Indeed, there is evidence to support that NPC quality control is compromised in ALS patients^54^. However, the studies addressing whether transport is altered in these C9ALS models have provided different answers, which can, at least in part, be explained by the differences in the methods used to study transport^29,47,51,52,55–58^ (reviewed in^40^). For example, it was shown that G_4_C_2_-repeats decrease importin α/β transport^29,47,51^, as do PR_20_ peptides^52^ and GR_25_^55^. In contrast, others only see an effect on importin α/β transport in the presence of GA_149_-GFP, and not with GFP-GR_149_ or PR_175_-GFP^56^, or determined none of the DPRs impacted importin α/β transport^57^. Also, in the presence of polyPR, the transport of the human NTR Transportin (TNPO1) was shown to be unchanged in some studies^56,57^, but reduced in another^51^.

Thus, while NCT and C9Orf72 ALS have been convincingly linked in literature, there are only few studies that directly address the changes in the kinetics of nuclear transport and the outcomes are not uniform nor comprehensive. Therefore, a clear answer to the question whether and how much the kinetics of nuclear transport of *all* proteins is impacted, has not been given. This is an important question because, if true, then a global derailment of protein localization should be expected in patients; a derailment that could only be solved at the level of interfering with the NCT machinery. However, it is also possible that the thus far reported connections are not a reflection of generally impaired nuclear transport kinetics, but rather a result of the mislocalization or aggregation of specific proteins that subsequently cause the mislocalization of a small subset of interacting proteins. Another outstanding question is whether transport defects are an effect of direct interaction of the disease-causing proteins with the nuclear transport machinery, or rather an indirect result from stress caused by these disease proteins. Here we take a reductionist approach and seek to answer these questions for simple models of polyPR or polyGA expressing yeast cells, measuring the changes in transport kinetics of four transport factors and comparing them to those measured under different stress conditions.

## Results

### Characterizing the components of the nuclear transport machinery in yeast C9ALS models

To study the effects of C9ALS DPRs on transport, we used a yeast model expressing 50 repeats of PR or GA, developed by^35^. 50 repeats is lower than common in ALS patients, but similar to previous studies (ranging between 10-65 repeats^35,38,39,48,49,51,55,59,60^) including a mouse model with 66 repeats which recapitulates molecular and behavioural abnormalities of C9ALS patients^61^. In accordance with previous studies^35–39^, the yeast cells expressing polyPR, but not those expressing polyGA, have a growth defect, as shown by the limited growth in a spot assay (Fig 1A). The reduced growth on plate by polyPR was validated by following individual dividing cells in microfluidic chips (ALCATRAS system as described^62^): we see that polyPR expressing cells have a median replicative lifespan of only two divisions, compared to a lifespan of eighteen cell divisions of WT cells (Fig 1B). In order to determine the localization of both DPRs, we tagged them (N-terminally) with mCerulean. This somewhat reduces the toxicity of polyPR, resulting in a median of six cell divisions (Fig 1B). mCerulean-polyPR is mostly nucleolar, since it co-localizes with the nucleolar protein Sik1, consistent with previous reports in other models^36–39,60^ (Fig 1C). PolyGA forms a rod-shaped focus in the cytoplasm (Fig 1C), which is also observed in other models^38,48,59,63–65^. Altogether, we conclude that our yeast model recapitulates basic features of toxicity and subcellular localization of the polyPR and polyGA DPRs as observed in other models.

**Fig 1.**
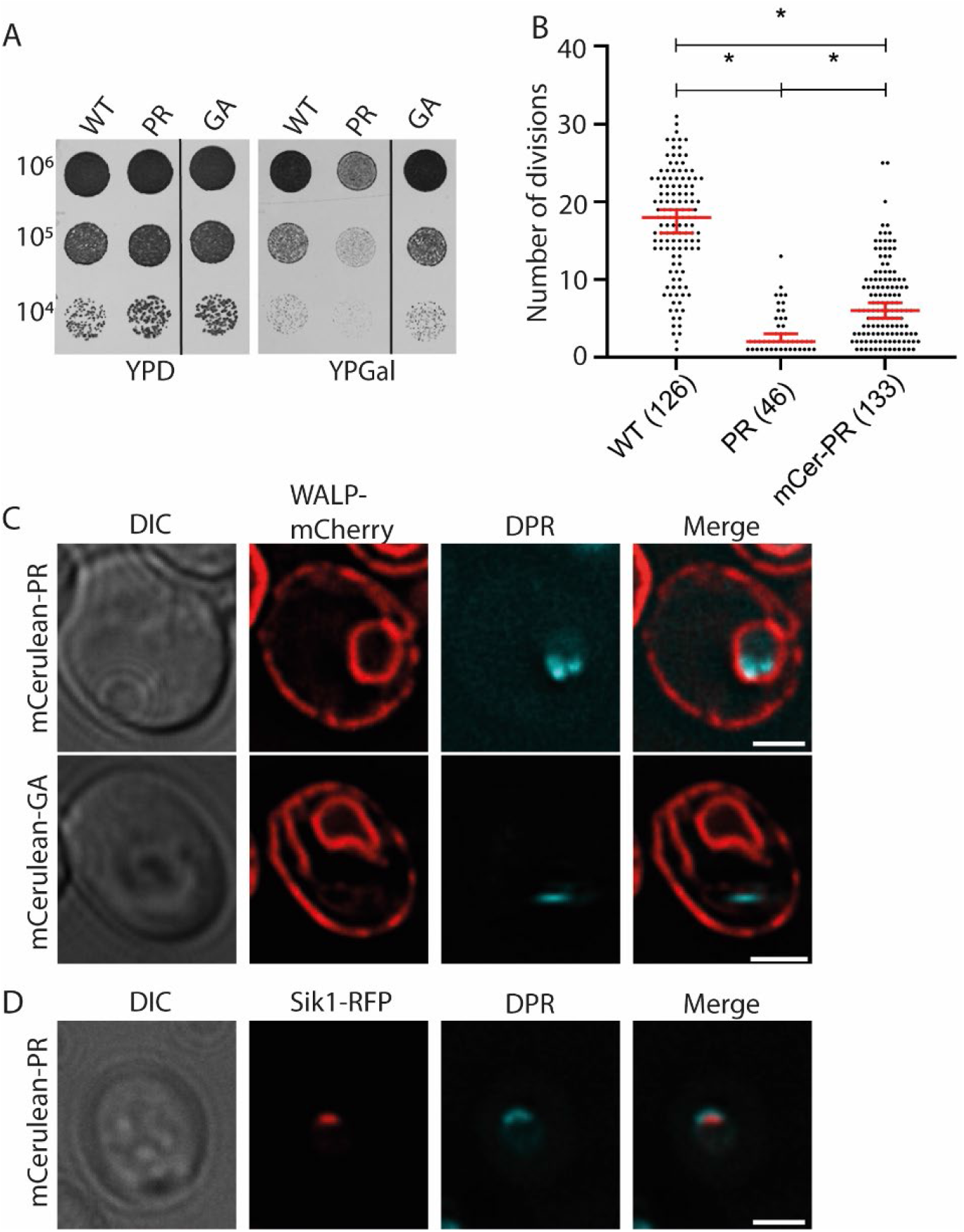
Yeast models for C9Orf72 ALS: expression of polyPR and poly-GA. **A)** PR_50_ is toxic under conditions when expression is induced while GA_50_ is not toxic. Left panel, uninduced (glucose), right panel, induced (galactose). **B)** Replicative lifespan of cells expressing PR_50_ or mCerulean-tagged PR_50_ (mCer-PR) in ALCATRAS chip compared to WT. Numbers of cells followed throughout their lifespan is indicated; data set from wild type cells taken from our previous work^72^. **C)** mCerulean-PR_50_ localizes in the nucleus, whereas mCerulean-GA_50_ localizes in a cytoplasmic focus. WALP-mCherry indicates the NE-ER network (in red). **D)** mCerulean-PR_50_ co-localizes with the nucleolar protein Sik1/Nop56-RFP, shown in red. Scale bar equals 2μm.

In order to determine whether the transport machinery components are altered in our model, we systematically studied the abundance and localization of the NTRs, the components of the NPC and those generating the Ran cycle (selection of NTRs and nucleoporins in Fig 2 and others in Fig 2 - supplement figures 1/2/3). First, we assessed the NTRs in our model. Alterations in the concentration of importing NTRs will change the transport kinetics, since the import rate of cargo is limited by the formation of NTR-cargo complexes in the cytosol^66–68^. Using SRM-based proteomics, we determined that the expression levels of the 18 NTRs found in yeast^5^ were similar in WT and polyPR or polyGA expressing cells (Fig 2 and Fig 2 – figure supplement 1A). To investigate the localization of these NTRs, we expressed PR_50_ or GA_50_ in strains expressing endogenously GFP-tagged NTRs, and were able to probe 16 out of the 18 yeast NTRs^5^ (Mtr2 and Ntf2 were unavailable). In general, we find no evidence that the localization of GFP-tagged NTRs is significantly altered upon expression of the DPRs. Small differences may be present in the localization of Pse1, which forms fewer nuclear envelope localized foci with both DPRs, and in the localization of Cse1, which is less abundant at the NE with PR_50_-expression (Fig 2 and Fig 2 – figure supplement 1B/C/D). Importantly, none of the NTRs accumulated visibly either in the nucleolus with polyPR, or in the cytoplasmic aggregate of polyGA. Jointly the analysis shows that expression of polyPR or polyGA does not inflict major changes in the abundance and localization of NTRs.

**Fig 2.**
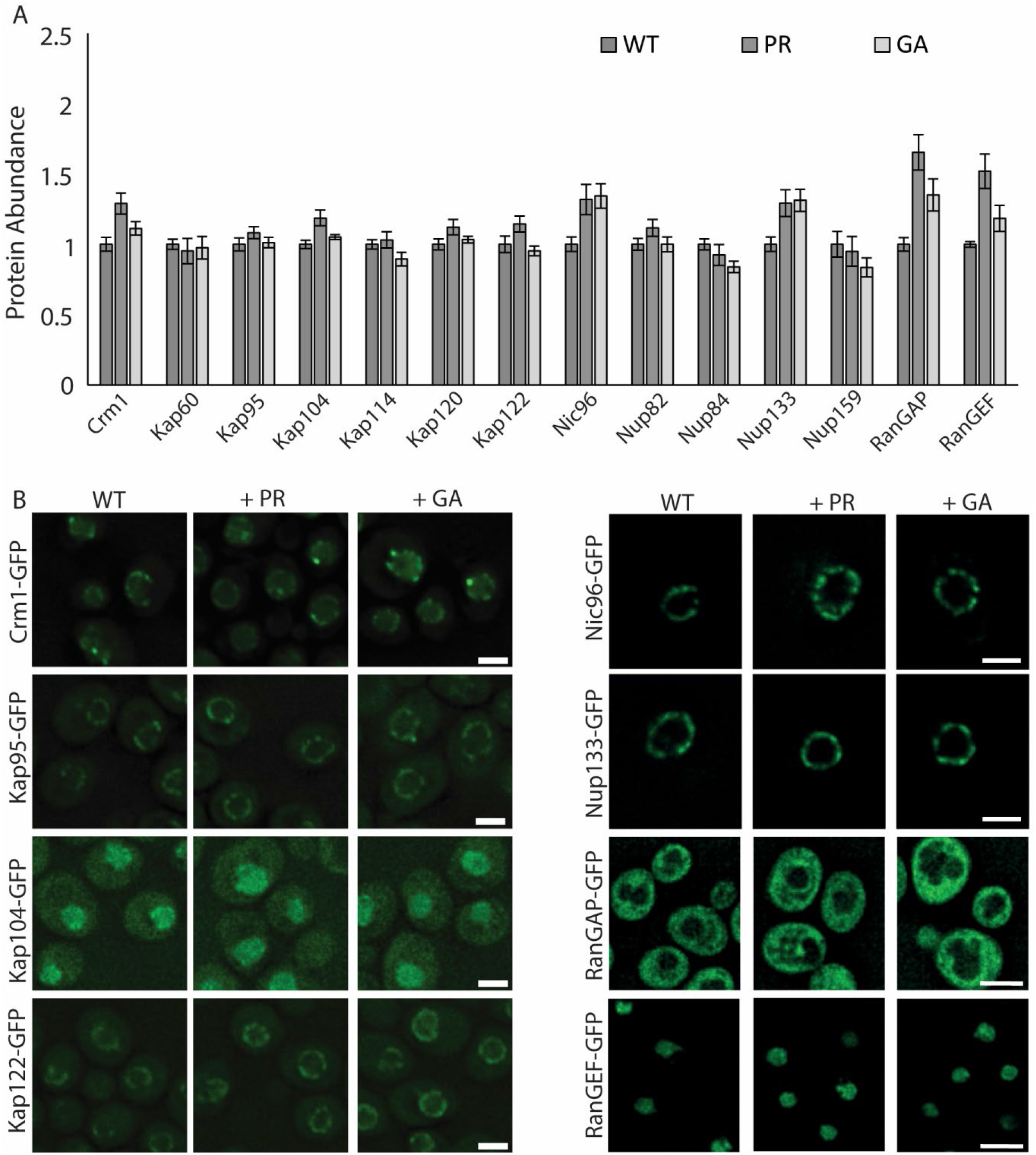
Abundance and localization of selected NTRs, Nups and the Ran system in polyPR and polyGA expressing cells. For other NTRs and Nups please see Figure Supplement 1,2,3. **A)** Abundance of few of the NTRs, Nups and RanGEF and RanGAP in whole cell extracts of WT or PR_50_/GA_50_-expressing cells determined by SRM-based proteomics in two biological and one technical replicate, source data 1. **B)** Localization of endogenously GFP-tagged NTRs compared between WT cells and cells expressing PR_50_ or GA_50_; scale bar equals 2μm.

Next, we assessed if there may be changes in the total number of NPCs in our ALS models. We measured the abundance of all nucleoporins in a proteomics analysis and found no expression level changes (Fig 2 and Fig 2 – figure supplement 2A). Also, the localization of four endogenously expressed GFP-tagged nucleoporins was assessed, and we see no relocation of the FG-Nups Nsp1 and Nup159, outer ring Nup133, nor the linker nucleoporin Nic96 (Fig 2 and Fig 2 – figure supplement 2B). Obviously, these measurements cannot report on the functional state of these NPCs, but the data does suggest that the expression of polyPR or polyGA does not majorly alter the numbers of NPCs.

Another major component of nucleocytoplasmic transport is the Ran cycle, which maintains the directionality of transport. Previous studies showed that the toxicity caused by (GGGGCC)_n_ RNA and poly-GR/PR is exacerbated when Ran and/or RanGAP1 are knocked down in *Drosophila* and yeast^35,47,50,69^. Co-localization of poly-GA aggregates with RanGAP1 has also been reported^48^, but this was not observed for Ran or RanGAP1 in another study^56^. In order to determine whether DPR expression changes the Ran cycle in our yeast model, we expressed endogenously tagged RanGAP-GFP, and its nuclear counterpart RanGEF-GFP, together with polyPR or polyGA. We did not observe any change in localization of either RanGAP or RanGEF (Fig 2 and Fig 2 – figure supplement 3A). Proteomics analysis shows that the total abundance of both RanGAP and RanGEF is possibly upregulated (a 1,5x fold increase is measured) in PR_50_ expressing cells, and unchanged in polyGA expressing cells (Fig 2 – figure supplement 3B). To assess if this change in RanGAP and RanGEF levels is associated with a change in energy levels, we measured cellular ATP levels with a FRET-based ATP-sensor (adapted from the AT1.03 ATP biosensor^70^). We find that the free ATP concentration is not altered in polyPR and polyGA expressing cells (Fig 2 and Fig 2 – figure supplement 3C). Since a depletion of ATP was shown previously to cause a drop in free GTP levels^71^, our measurement of stable ATP levels may indicate that also GTP levels are unchanged in polyPR and polyGA expressing cells.

In summary, our yeast model recapitulates the toxicity and localization of the DPRs polyPR and polyGA as reported in other models. Our analysis shows that the abundance and localization of NTRs, nucleoporins, and thus also NPCs, are unaltered upon expression of polyPR or polyGA. In PR_50_ expressing cells a small upregulation of RanGAP and RanGEF was measured, but at least energy levels as measured via free ATP levels are unchanged. Having characterized the components of the nuclear transport machinery in the yeast ALS models, we next study whether transport is affected by either polyPR or polyGA expression.

### Does expression of polyPR and polyGA change import and export?

Having characterized the abundance and localization of the main players in NCT, we next determined the effect of DPRs on two functional read-outs for nuclear transport^73,74^, namely the rate of passive diffusion and the rate of active transport through the pores. For mobile proteins, the balance between NTR-facilitated import of NLS-containing proteins and their passive efflux leads to nuclear accumulation. Similarly, the balance of NTR-facilitated export of proteins with an NES and their passive influx leads to a steady-state nuclear exclusion (Fig 3A). Not all NTRs and their NLS/NESs are known, and in this study we have used i) the Stress-Seventy subfamily B1 NES (NES_Ssb1_^75^) for the Crm1 exportin, ii) the monopartite, classical Simian Virus 40 NLS (NLS_Sv40_^76^) for the Kap60/Kap95 import complex, iii) the bipartite nucleophosmin (Npm1) NLS (NLS_Npm1_^77^) also for the Kap60/Kap95 import complex, iv) the Nuclear polyAdenylated RNA-Binding 2 NLS (NLS_Nab2_^78^) for Kap104, and v) the PHOsphate metabolism 4 NLS (NLS_Pho4_ ^79^) for Kap121/Pse1 (Table 1 and more details in Table S1).

**Fig 3.**
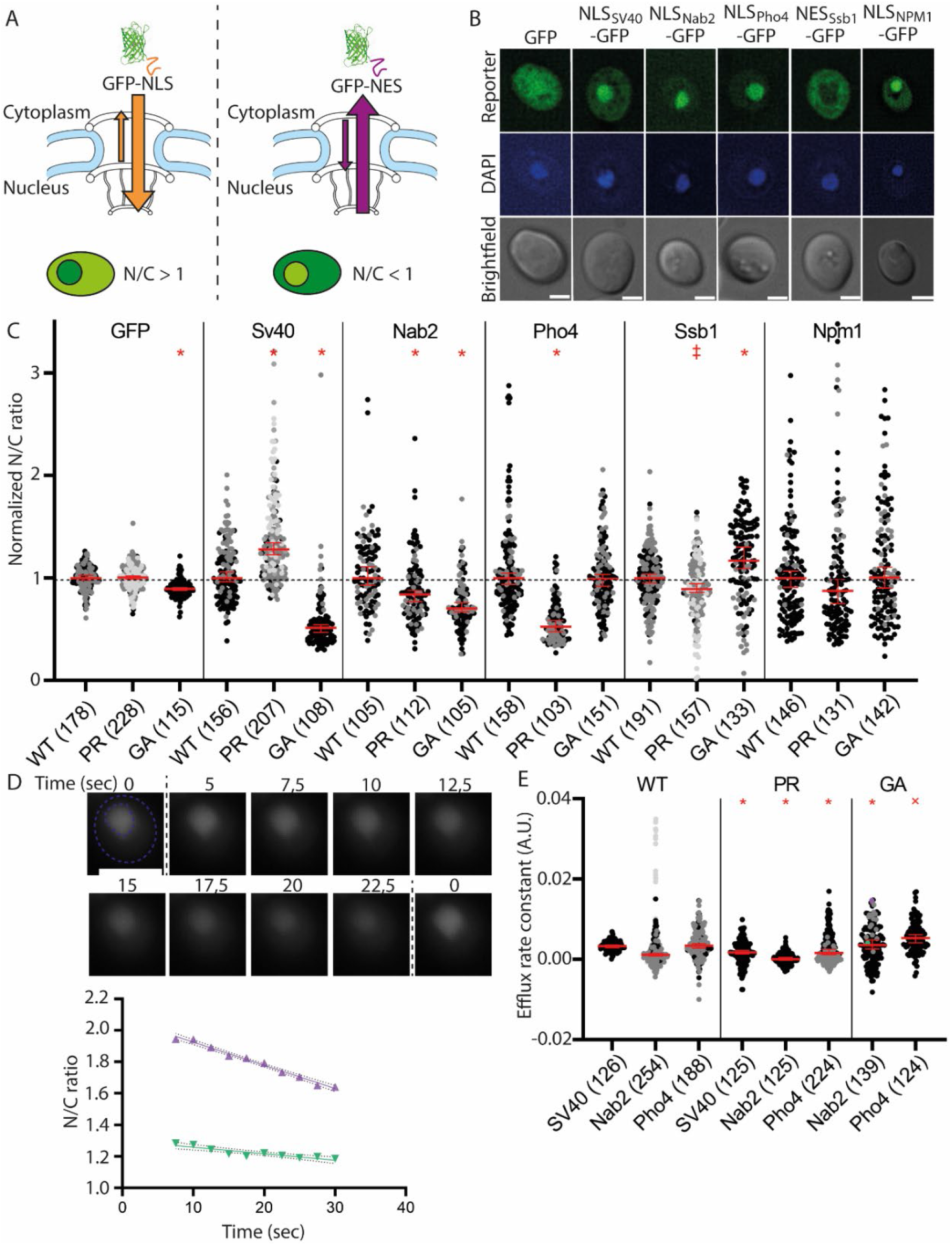
Steady-state N/C ratios of transport reporters are altered with polyPR and polyGA expression. **A)** Nuclear accumulation or exclusion are the result of active import and passive leakage resulting in nuclear accumulation for the NLS-reporter, and the opposite for the NES-reporter. **B)** Subcellular distribution of the GFP-reporters in wildtype cells, nuclear staining DAPI, scale bare equals 2μm. **C)** N/C ratios of GFP-transport reporters in WT, PR_50_ or GA_50_-expressing cells, determined in 2-4 independent experiments (grey colours); total number of cells analysed between brackets, source data 3. Data is normalized to the median WT values for each reporter, median and 95% confidence interval are shown in red, Mann-Whitney comparison to WT with ‡ = p-value <0.01, and * = p-value <0.0001. A total of 10 datapoints with normalized N/C above 3.5 are not shown but taken into account in the statistics (respectively 1,1,1,2 and 5 points for the datasets *SV40-PR, Pho4-WT, Pho4-GA, NPM1-PR* and *NPM1-GA*). **D)** Measurements of transport reporters nuclear accumulation loss and calculated efflux kinetics. Time zero is the time point at which the metabolic poisons (Na-azide and 2-deoxy-glucose) reached the cells and active transport of the reporters stops. N/C ratios were recorded 5-22,5 sec after poisoning occurred, after which linear regression slopes were determined. Green triangles show a cell with average efflux rate constant of -0,0041; purple triangles show the presented cell with a particularly strong reduction of the N/C ratio of -0,014. **E)** Efflux rate constants in WT and ployPR and polyGA expressing cells, data from independent experiments are shown in different grey colours, with the median and 95% confidence interval in red, Mann-Whitney comparison to WT with × = p-value <0.001, and * = p-value <0.0001.

**Table 1.**
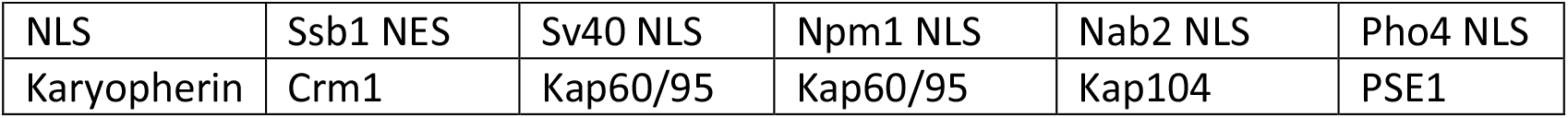
NLSs and NTRs

| Table 1 NLSs and NTRs |  |  |  |  |  |
| --- | --- | --- | --- | --- | --- |
| NLS | Ssb1 NES | Sv40 NLS | Npm1 NLS | Nab2 NLS | Pho4 NLS |
| Karyopherin | Crm1 | Kap60/95 | Kap60/95 | Kap104 | PSE1 |

To monitor the effect of PR_50_ or GA_50_ on nuclear transport, we quantified the steady state localization of GFP-NLS reporter proteins encoding the NLSs specific to Kap60/95, Kap104, and Pse1, and a GFP-NES reporter recognized by Crm1 (summarized in table S1, and Fig 3B) in cells expressing PR_50_ or GA_50_. As a control for the reporter proteins, we included GFP without sorting signals. The steady state distribution of the GFP-reporters is calculated by taking the ratio of the fluorescence measured in the nucleus and the cytosol (the N/C ratio). GFP accumulates slightly in the nucleus due to its hydrophobic surface properties^80^, represented by a median N/C ratio of 1.24 ± 0.14 in WT cells (Fig 3C, and table S2). The N/C ratio of GFP is identical in cells expressing PR_50_, and slightly lower in GA_50_ expressing cells (median N/C ratio to 1.11 ± 0.11). The import reporters accumulate in the nucleus, and thus have an N/C ratio >1, whereas the export reporter is excluded from the nucleus and has an N/C ratio <1 (Fig 3B). Please note that when nuclear export decreases, the N/C ratio will increase and approach the value of 1. We interpret a decrease in steady state nuclear localization of the import reporter as a decrease in the import kinetics by this NTR. This interpretation holds under the assumption that the import of native cargo’s (that compete with the reporter for the same pool of NTR^81^) is stable under the conditions we compared. Support for this comes from proteomics data showing that the abundance of a few known cargoes of the NTRs involved (Kap95, Kap104, Pse1 and Crm1) are not altered upon DPR expression (Source data 2). In addition, our data (Fig S1A) and previous studies following the transport of GFP-NLS reporter proteins^67^, suggest that the NTR availability is not limiting under the conditions used.

We first addressed if expression of the DPRs alters the passive permeability of the pore. For this we measured the immediate loss of nuclear accumulation of import reporters carrying the Sv40, Nab2 or Pho4 NLS after inhibiting active transport (Fig 3D). The inhibition of active transport is achieved by poisoning the cells with sodium azide and 2-deoxy-D-glucose, which respectively inhibit the mitochondrial respiratory chain and glycolysis. The analysis showed that the expression of PR or GA resulted in minimal changes in leak rate for the NLS_Sv40_-, NLS_Nab2_- and NLS_Pho4_-GFP reporter (Fig 3E, table S3).

Having established that polyPR or polyGA expression gives only minor changes in passive permeability of NPCs, we measure the steady state localization of import and export reporters to probe also the active transport rates. Assessing the effects of polyPR expression we see that the nuclear exclusion of NES_Ssb1_-GFP and the nuclear accumulation of NLS_Sv40_-GFP are both modestly but significantly increased when PR_50_ is expressed. However, the NLS_NPM1_ reporter is not significantly affected by polyPR expression, showing that even for one transport route, i.e. via the Kap60/95 complex, different effects are observed for specific NLSs. In contrast, a loss of compartmentalization is observed for the reporters carrying the NLS_Nab2_ or the NLS_Pho4_. The effects on transport caused by polyGA expression are distinct: export is reduced in polyGA expressing cells, and import of the NLS_Sv40_-GFP and NLS_Nab2_-GFP are also decreased (Fig 3C and table S2). The nuclear accumulation of both the NLS_Pho4_-GFP and the NLS_NPM1_-GFP reporter is unaffected. It is interesting that for both polyPR and polyGA expression, despite the variation observed in the other transport routes, the NPM1 reporter (carrying a bipartite NLS) is unaffected.

In short, the transport measurements jointly show that polyPR increases the transport of the classical NLS_SV40_, and the NES_Ssb1_, but decreases transport of the other two import routes. PolyGA reduces transport of the NLS_Sv40_, NLS_Nab2_, and NES_Ssb1_. Thus, the analysis highlights that DPR expression impacts the transport facilitated by different NTRs, and even by NLSs for the same NTR, and that to varying degree and direction.

### Nuclear transport under stress conditions

So far, our analysis established that DPR expression impacts the transport kinetics by different NTRs, but it is not clear if we should consider these changes to be small or large. To provide a context for the quantitative assessment of transport defects, we compare them to those measured under several stress conditions. We subjected yeast to diverse stressors, namely heat shock, starvation, osmotic shock, and oxidative stress, and determined the effect on transport.

Upon mild heat shock at 37°C for 5 minutes, classical import (NLS_Sv40_) and classical export (NES_Ssb1_) are not changed, but import of the NLS_Nab2_- and NLS_Pho4_-GFP reporters is reduced (Fig 4B, table S4). If cells are kept at mild heat shock for an hour, they adapt to the stress and the import by all three importins goes up again, even increasing above wild type levels. This overshoot effect is the strongest for the NLS_Pho4_-GFP reporter. Interestingly, increasing the heat shock to severe temperatures – i.e. 42°C for 5 minutes – shows decreased import of the NLS_Sv40_-reporter, but now the NLS_Pho4_-reporter is not affected. Under sublethal heat shock at 46°C for 10min the differences between the import routes become even larger, where import of the NLS_SV40_-reporter is further reduced compared to 42°C, import of the NLS_Nab2_-reporter is increased, and the NLS_Pho4_-reporter is not affected. Also interesting is that the NLS_Pho4_-reporter is only impacted at 37°C, but under higher temperatures import is not affected. Next, we subjected the cells to two different methods of starvation. When cells pass the diauxic shift, after growing to saturation, we see a decrease in import of the NLS_Sv40_-reporter, and a decrease in export of the NES_Ssb1_-reporter after 20 hours in medium (Fig 4C and table S4). The transport of the other two import reporters is not significantly decreased. In contrast, after 24h in water, all three import routes, as well as export are reduced. Thus, import by these three pathways is more susceptible to water shift, whereas the impact of saturated growth is stronger on export.

**Fig 4.**
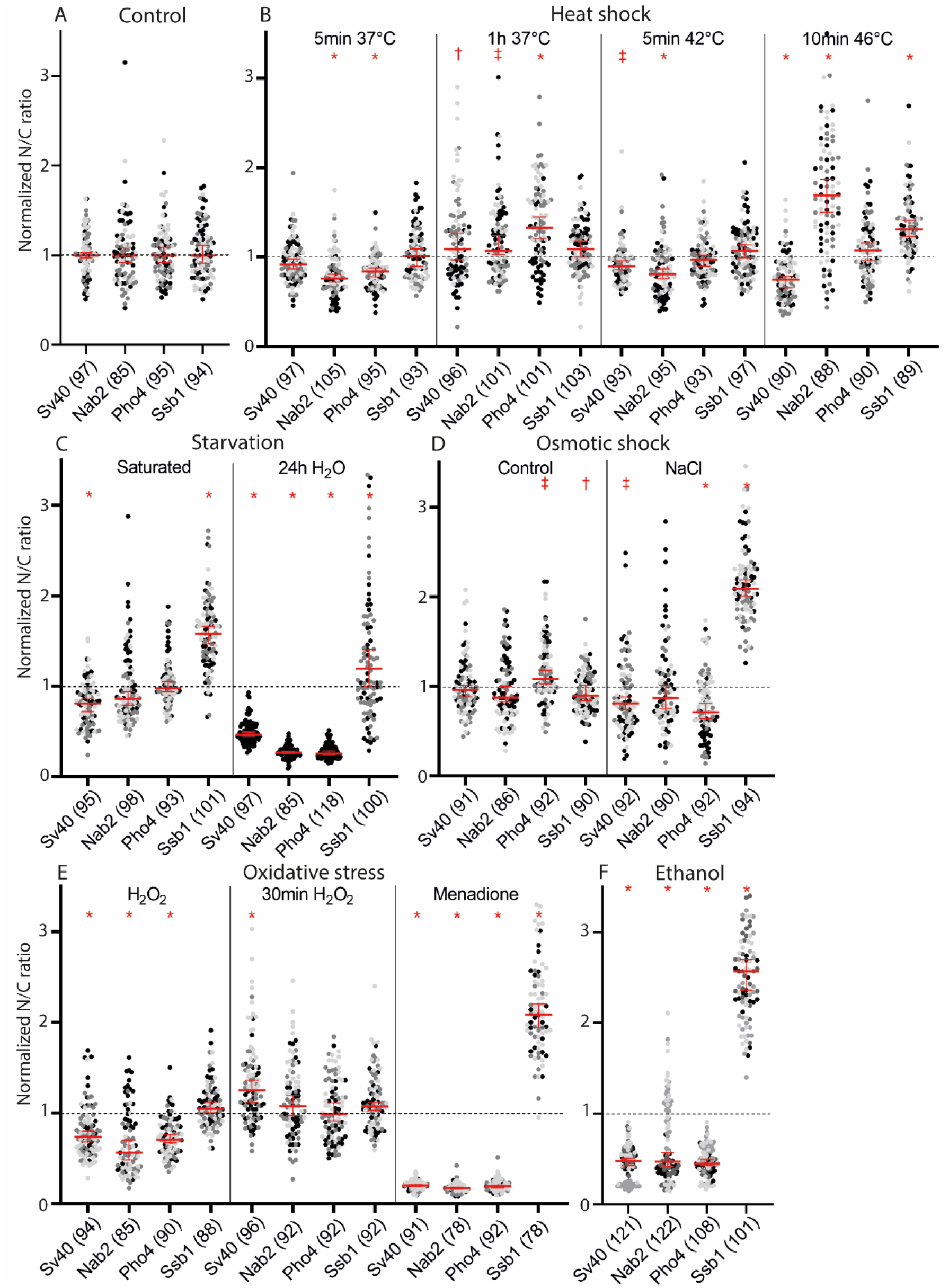
Steady-state N/C ratios of transport reporters under environmental stress conditions. N/C ratios of GFP-transport reporters in control (A) and indicated stress conditions (B-F), determined in 2-4 independent experiments (grey colours); total number of cells analysed is indicated between brackets, source data 4. Data is normalized to the median control values for each reporter, median and 95% confidence interval are shown in red, Mann-Whitney comparison of each reporter to WT with † = p-value <0.05, ‡ = p-value <0.01, * = p-value <0.0001. A total of 31 datapoints with normalized N/C above 3,5 are not shown but taken into account in the statistics (respectively 3, 10, 2, 1, 1, 14 points for the datasets in B (Nab2-1h37C, Nab2-10min 46C), C (Ssb1-24h), D (Nab2-NaCl, Ssb1-NaCl), E (Ssb1-Menadione) and F (Ssb1-ethanol). **A)** Wild type BY4741 cells expressing the transport reporters. **B)** Heat shock was performed at 5 minutes or 1 hour at mild (37°C), 5min at severe (42°C), and 10min at sublethal (46°C) temperature stress. **C)** Starvation was performed by growing the yeast cultures to saturation in 20h, or replacing the medium with water. **D)** Osmotic shock was measured immediately after adding 1M NaCl to cells adjusted to the NaPi buffer as control. Normalization of this category is to the osmotic control instead of WT. **E)** Oxidative stress was induced with 4mM H_2_O_2_, and measured immediately or after 30minutes, or by adding 1,2mM Menadione for 40min. **F)** Ethanol stress was performed by adding 15% ethanol to the cultures for 20 minutes.

Next we assessed the response to an osmotic shock with 1M NaCl (as in^82^) (Fig 4D, table S4). We see that both the NLS_Sv40_- and NLS_Pho4_-reporter are less accumulated under osmotic shock. The NLS_Nab2_-reporter is not significantly impacted, and the export of the NES-reporter is drastically impaired.

A last set of stresses induces oxidative stress, either by means of the addition of 4mM hydrogen peroxide, or via the intracellular production of reactive oxygen species (ROS) after 40 minutes exposure to 1,2mM menadione. Peroxide stress reduces import of all three different reporters, but export is not affected (Fig 4E, table S4). Hydrogen peroxide, being highly reactive, will only temporarily generate ROS, and indeed 30 minutes after the addition of hydrogen peroxide the cells return to normal transport phenotypes. Similar to what we saw with adaptation to mild heat stress, classical import “overshoots” the normal accumulation and even increases import a little over the wild type situation. ROS created by incubation with menadione completely abolished active transport: the N/C ratios of the import reporters are all reduced to roughly 1. Export is drastically reduced, similarly to osmotic stress, but the N/C ratios are still below 1.

Lastly, incubation of the cells with 15% ethanol for 20 minutes reduced import strongly, although it is not abolished and N/C ratios are still above 2 (Fig 4F, table S4). Export is completely inhibited, and the N/C ratio reaches 1.

In conclusion, we have quantified transport by four NTRs under twelve different environmental stressors. Our findings align with previous work in yeast^18^ measuring the transport by one NTR (Kap60/95) in stationary phase culture, and in conditions of exposure to ethanol, oxidative stress, heat stress, and adaptation to heat stress. They are different for those measured in condition of osmotic shock; possibly due to differences in the model^18^.

Next, we aimed to compare the DPR induced transport defects to those induced by making genetic alterations of components of the nucleocytoplasmic transport machinery. First, we used knockouts of five different non-essential NTRs, each one not required for the transport of any of our transport reporters, namely Sxm1Δ, Kap114Δ, Kap120Δ, Kap122Δ, and Kap123Δ. These five NTRs have different expression levels, cargoes, and localizations (Fig S1B). In general, we find that the knockout of an NTR leads to a greater variability in calculated N/C values of our transport reporters. This is also observed with the environmental stresses and DPR expression, and is possibly a reflection of a derailed cell physiology (compare Fig 4A and Fig 5A, raw values in table S5). Excitingly, the NTR knockouts increase the import of the NLS_Sv40_-reporter, similar to what we saw with polyPR expression. The import of the NLS_Nab2_- and NLS_Pho4_-reporters was decreased in four of the knockout strains, but not in the Kap114Δ. Kap120Δ increases the import of all three NLS-based reporters. Export in the Sxm1Δ, Kap120Δ, and Kap122Δ strains was decreased, but not significantly altered in Kap123Δ. As a second genetic alteration, we overexpressed either RanGAP or RanGEF to see how a misbalance in the Ran cycle would impact transport. We had anticipated that the effect of both overexpressions would be a decrease in transport, since both result in an alteration of the RanGTP gradient. The overexpression of RanGAP and RanGEF does indeed decrease the import and the export of our reporters, but the extent of the effect is variable for the different reporters (Fig 5B, table S5). The NLS_Nab2_-reporter is more susceptible to the overexpression of RanGAP, while export is only impacted when RanGEF is overexpressed.

**Fig 5.**
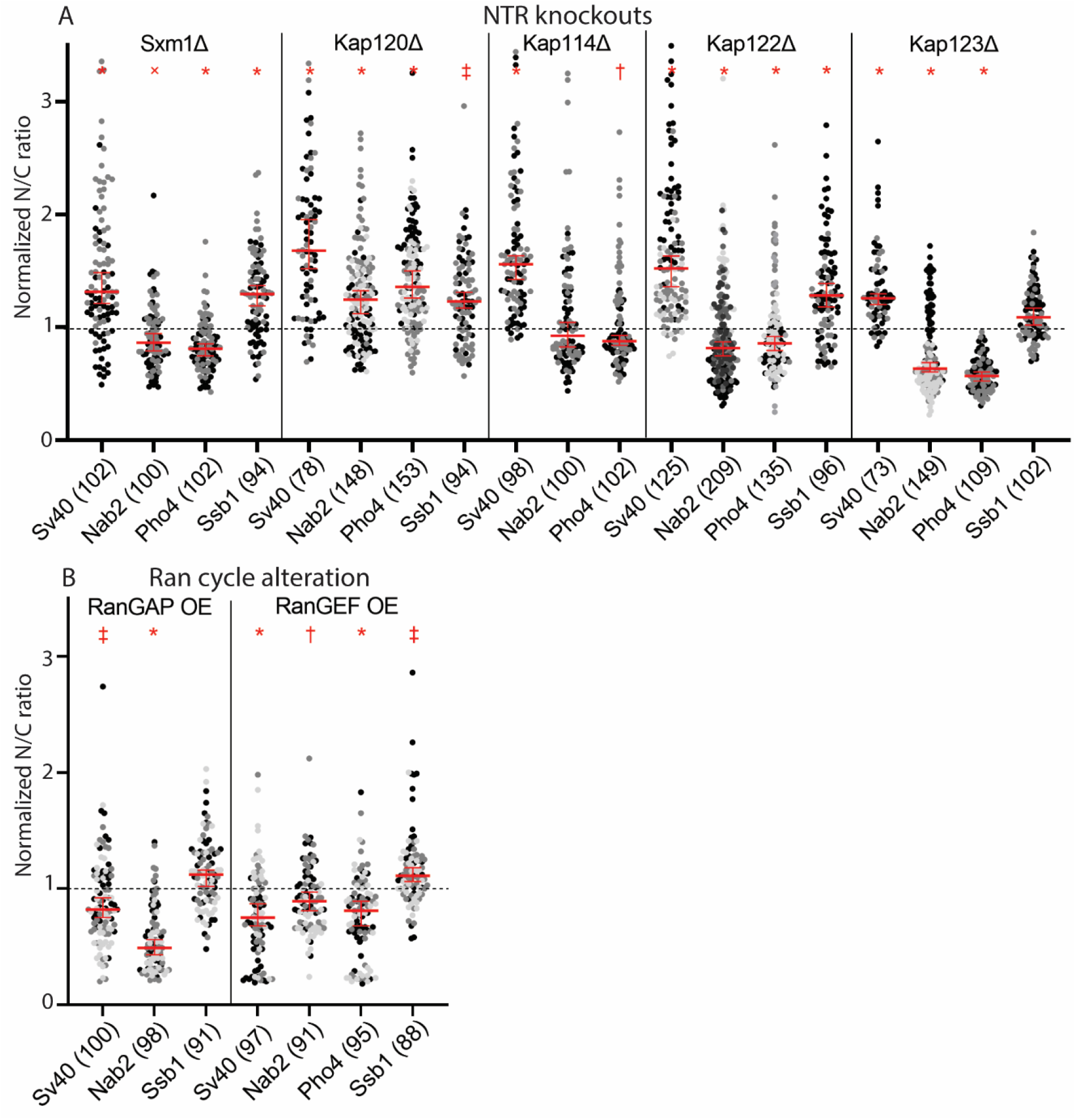
Steady-state N/C ratios of transport reporters in NTR and Ran-cycle mutants. **A)** N/C ratios of GFP-transport reporters in WT, and NTR knockout cells, determined in 2-4 independent experiments (grey colours); total number of cells analysed is indicated between brackets, source data 5. Data is normalized to the median WT values for each reporter, median and 95% confidence interval are shown in red, Mann-Whitney comparison of each reporter to WT with † = p-value <0.05, ‡ = p-value <0.01, × = p-value <0.001, * = p-value <0.0001. A total of 8 datapoints with normalized N/C above 3,5 are not shown but taken into account in the statistics (respectively 1,2,2,1,1,1 and 5 points for the datasets SV40-Sxm1Δ, Nab2-Kap120Δ, Pho4-Kap120Δ, Nab2-Kap114Δ, SV40-Kap122Δ, Nab2-Kap122Δ). **B)** As in A but now for cells overexpressing RanGAP or RanGEF.

In general, we see that the transport of Nab2-containing cargo, as performed by Kap104, is the most variable under the different conditions probed, and an outlier under sublethal heat stress (Fig 4). Export is in general the most stable, although remarkably susceptible to osmotic shock. A comparison between the sizes and directions of the effects on transport induced by the different environmental stressors, mutations and DRPs, shows that they provide unique profiles but, with the exception of the more extreme interventions (24h in water, menadione, ethanol), the effects are of comparable size. The extent of derailment of nuclear transport induced by DPRs is thus to be expected in the same size range as those induced by natural stress conditions such as starvation or osmotic stress. Comparing the NCT phenotypes under these stresses to the data on NCT in polyPR expressing cell, it stands out that whereas most stresses reduce transport, albeit to different extents, NTR knockout leads to an increase in NLS_Sv40_-reporter import, which matches with the effect polyPR expression has.

### Similarities between transport effects in C9-ALS models and stress conditions

To further investigate which stress-induced transport phenotype shows similarity with the transport phenotypes observed in the C9ALS model, we performed clustering analysis of the transport datasets. We first performed a Principal Component Analysis to show how similar or dissimilar the stresses are, and which component (i.e. transport reporter) mainly influences this (dis)similarity. This is represented by a biplot (Fig 6A), which shows that the phenotypes for the strong effectors menadione, ethanol, and water shock, are clearly very different from the other stresses. It does not allow analysing the similarities and differences in the conditions inducing more subtle changes. However, when clustered in 5 categories, as by the k-means clustering analysis, we find more meaningful clustering, as e.g. now the conditions that increase classical transport become a separate category. These categories are represented in the biplot with different colours of the labels and in the heatmap in separate blocks (Fig 6A/B). The first cluster includes menadione induced ROS and is characterized by a strong defect for the import reporters reducing the N/C ratios to 1 (Fig 6B & table S4). The second cluster of water starvation and ethanol includes basically the two ‘outliers’ in the biplot, where the common factor is the reduction of the classical import route, but to a smaller degree than we saw for menadione treatment. The next three categories are the biologically more relevant ones. Cluster 3 includes PR expression, and the NTR knockout strains Sxm1Δ, Kap122Δ, and Kap123Δ; cluster 4 includes Kap120Δ, the osmotic shock control, peroxide after 30min, mild heat shock after 1h, and 10min sublethal 46°C heat shock, and cluster 5 includes peroxide stress, phosphate buffer, 5min mild and severe heat shock (37°C and 42°C), RanGEF overexpression, GA expression, saturated growth and osmotic shock. Kap114Δ and RanGAP OE were not included in the PCA, because one category of transport data was lacking. However, the similarities in the pattern of the effects, as shown in the heatmap and the bar graphs, allowed us to manually add Kap114Δ with the other NTR knockouts in group 3, and RanGAP OE with group 5.

**Fig 6.**
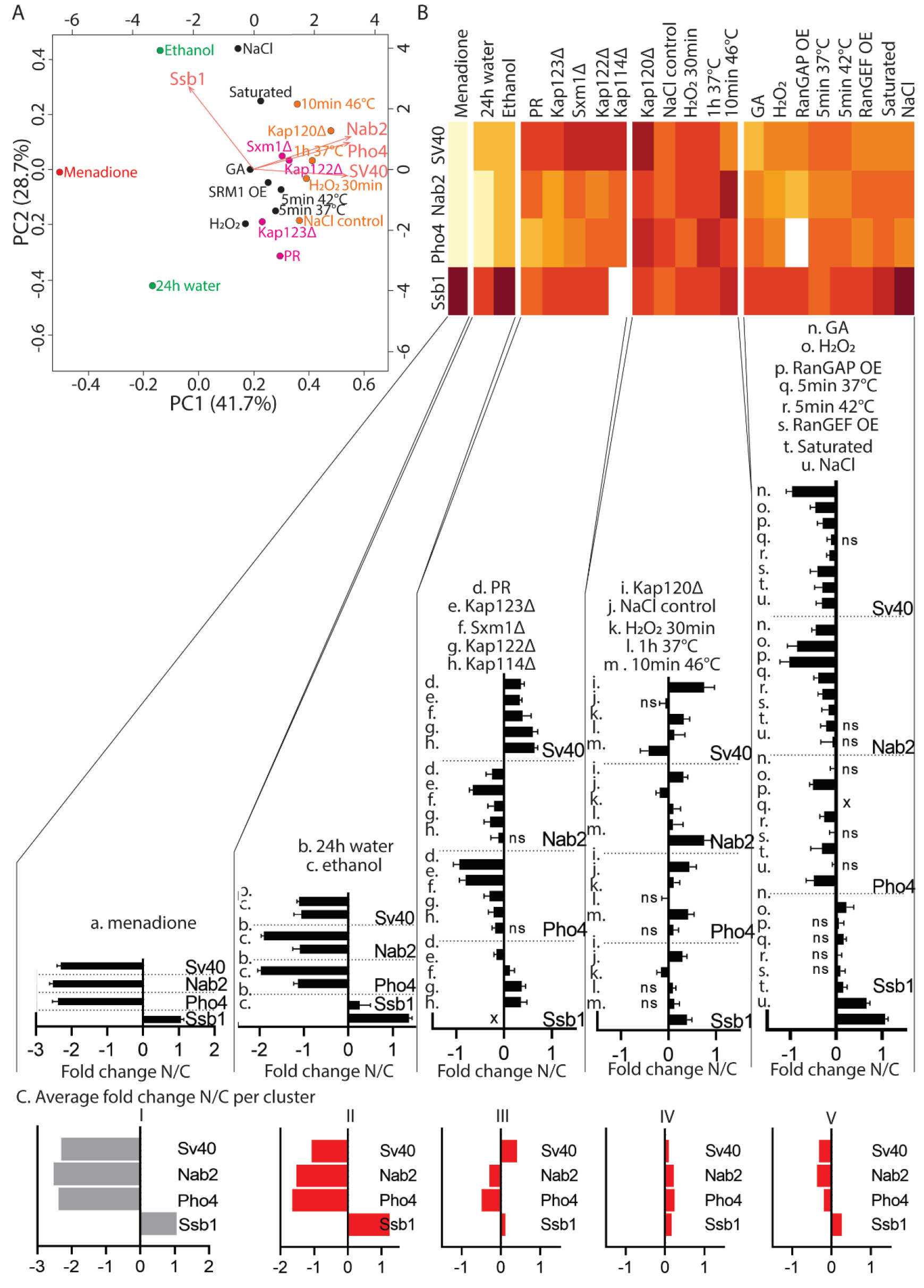
Similarities and differences in nuclear transport defects. **A)** K-means clustering leads to five categories, shown here on a biplot based on the median values of the four transport reporters, each group in one colour. **B)** The five categories as shown by their fold change for the different transport reporters. Cluster 1) the strongest change is seen for menadione stress and 24h water stress; these strongly reduce import and export, menadione reduces import to the maximal extent of reaching N/C ratios of 1. Cluster 2) 0.5h water and ethanol stress strongly reduce export, with ethanol reducing export to N/C ratios of 1, and reduce import less strong than group 1. Cluster 3) PR expression, and the NTR knockout strains Kap123Δ, Sxm1Δ, Kap122Δ, and Kap114Δ show an increase in import of the SV40 reporter, and reduced import of the other reporters. Export is reduced for the NTR knockouts, but increases for the PR expressing strain. Cluster 4) Kap120Δ, the osmotic shock control, peroxide after 30min, mild heat shock after 1h, and 10min sublethal 46°C heat shock are grouped based on a slight increase in import, and small decrease in export. Cluster 5) peroxide stress, RanGAP overexpression, phosphate buffer, 5min mild and severe heat shock (37°C and 42°C), RanGEF overexpression, GA expression, saturated growth and osmotic shock are categorized by a general decrease of import, while export is unaltered or reduced. **C)** Shows the average N/C fold change of the intervention in the cluster.

We will describe the characteristics of cluster 3-5 in more detail: Cluster three is dictated by the improved import of the Sv40-reporter, together with reduced Nab2- and Pho4-reporter import. The effects on export are dissimilar, as they are increased in polyPR-expressing cells, and reduced in the NTR knockout strains. The average pattern of changes in transport is shown in figure 6C.

The fourth cluster gathers adaptations to environmental stresses and is mainly determined by the increase of Nab2- and Pho4-reporter import, and in part by the increase of Sv40-reporter import and reduced export. Kap120Δ shows the strongest improvement of import. The conditions probed reflect adapted cell states where the transport returns to wild type situation. Cells placed in NaPi buffer show reduced import in the first ∼10minutes, but later adapt to this buffer (NaPi shock compared to osmotic control in table S4). Also, peroxide initially triggers a response (see cluster 5) but peroxide readily degrades and is no longer present after 30 minutes, when cells have adapted.

The fifth cluster has the smallest changes in NCT and is defined by reduced import, but less strong than in the first two clusters, as well as reduced export. The common denominator for this category is the alteration of energy maintenance conditions: RanGAP and RanGEF overexpression affect the Ran cycle directly; peroxide stress impacts mitochondria; saturated growth limits glucose availability and imposes mild oxidative stress linking back to mitochondrial stress; the heat shock is related to energy generation^83^ and the heat shock response regulates glycolysis as one of the key processes^84^; and osmotic shock is related to metabolic changes^85^.

In short, we see that different stresses have variable effects on transport, and we can distinguish categories which overlap with the effect or polyPR and polyGA expression. In this case, it allows us to cluster polyPR expression with alterations in NTR levels, and polyGA expression with stresses that affect energy maintenance. If we consider that changes in NCT for the four import routes and the export route may serve as a fingerprint reporting on cell physiology, then we may infer that polyPR and polyGA expression impact the maintenance of, respectively, NTR and energy levels.

Here we specifically address a possible cellular mechanism (that has also previously been proposed^51,55^), namely, that polyPR binds to NTRs which decreases their availability for transport. In cells, NTR levels are finely tuned, the majority of NTR knockouts are inviable, and overexpression of NTRs leads to toxicity (Fig S2). The distinct behaviour of Kap120Δ provided an entry point into the question whether polyPR may alter NCT by binding NTRs. As shown in figure 5A, the transport profile of Kap120Δ is different from the other four: while Sxm1Δ, Kap114Δ, Kap122Δ, and Kap123Δ show respectively an increase, a decrease, a decrease and an increase in the N/C ratio reporting on the transport of NLS_Sv40_-, NLS_Nab2_, NLS_Pho4_, and NES_Ssb1_-GFP reporters, in Kap120Δ cells all reporters have an increased N/C ratio. Previously it was shown that polyPR can bind importins^51,55^ and hence we investigated the possibility that the distinct behaviour of transport in Nup120Δ cells is related to a difference in its capacity to bind polyPR.

To compare the capacity of the NTRs Kap120 and Kap114 to bind to polyPR, we performed coarse-grained molecular dynamics simulations. For this purpose, we developed residue-scale models of Kap120 and Kap114 for which the crystal structures of unbound state are known (PDB codes 6fvb and 6aho, respectively). The coarse-grained models preserve the overall structure of Kap120 and Kap114 (Fig 7A). Additionally, the distribution of charged and aromatic residues that are relevant for the interaction of arginine-rich DPRs with NTRs are included in the model. For polyPR, we used a coarse-grained model that has already been applied to study the phase separation of polyPR with negatively-charged proteins^86^. More details about the coarse-grained force field are provided in the Materials and Methods section. We observed direct binding of Kap120 and Kap114 to PR_50_ at monovalent salt concentrations of 200mM and 100mM. To quantify the binding, we calculated the time-averaged number of contacts C_t_ between polyPR and the NTRs (Fig 7B). As can be seen, reduction of the salt concentration, which reduces screening effects, increases the number of contacts, thus showing the important role of electrostatic interactions in polyPR-NTR binding. For C_salt_ = 200mM, a condition similar to previous *in vitro* experiments^55^, we observe a significantly lower number of contacts (around 2 times lower) for Kap120 compared to Kap114. This difference in binding behavior can be related to a difference in the net charge of these two NTRs.

**Fig 7.**
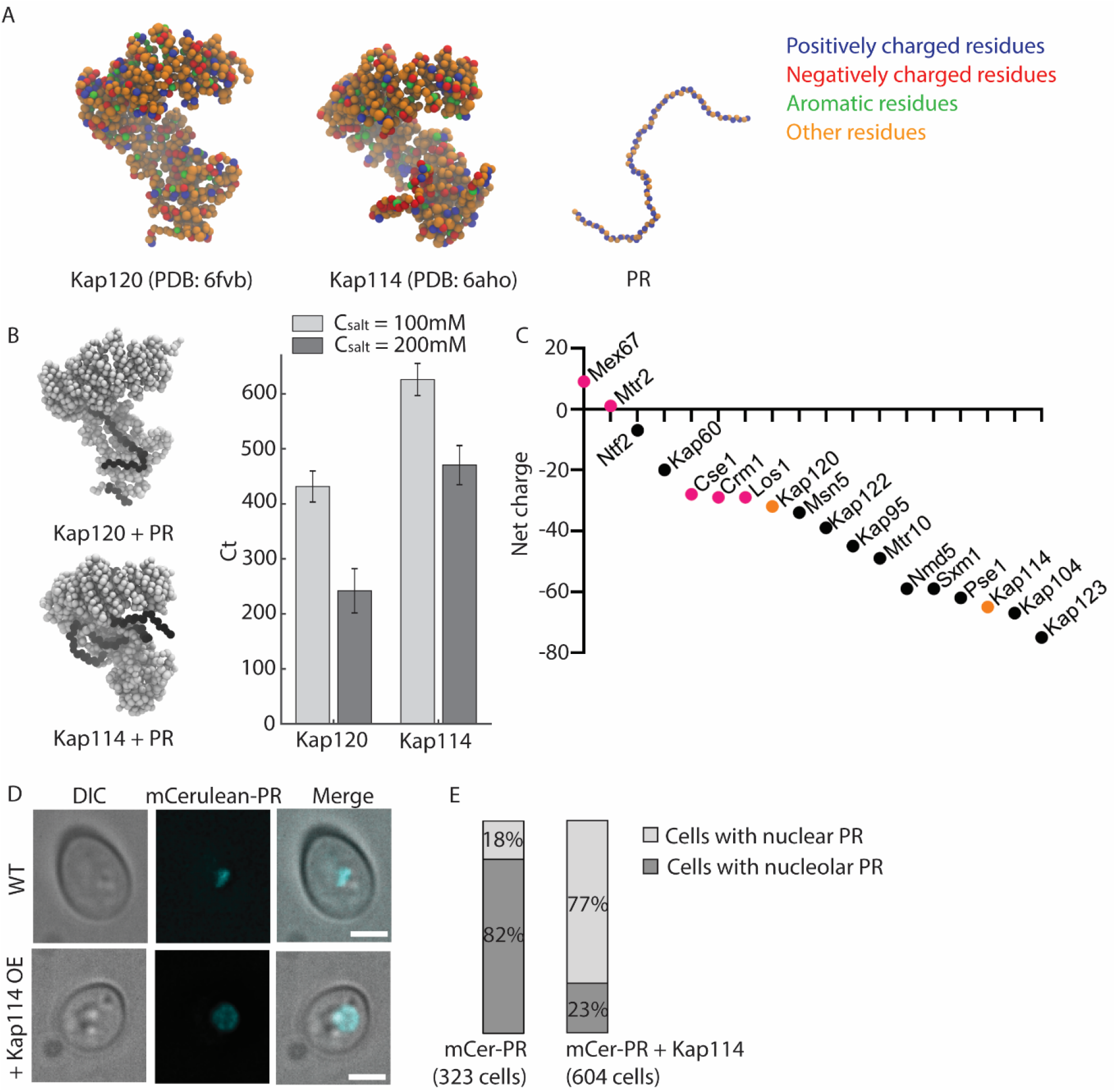
Investigating electrostatic interactions of Kap114 and Kap120 with polyPR. **A)** Schematic depiction of residue-scale coarse-grained models of Kap120, Kap114, and PR_50_ with the different bead types highlighted. **B)** PR_50_ time-averaged number of contacts Ct with Kap120 and Kap114 calculated at two different monovalent salt concentrations C_salt_ = 100mM and 200mM. Two snapshots for PR_50_-Kap120 and PR_50_-Kap114 binding at C_salt_ = 200mM are shown on the left, with polyPR depicted in black and NTRs in grey. Simulations are performed for 2.5μs and the last 2μs are used for the contact analysis. **C)** Overview of the net charge of each NTR, by subtracting the positively charged residues (Arg/Lys) from the negatively charged residues (Asn/Glu), source data 6. Colour-coded are the exportins in magenta, and the Kap120 and Kap114 in orange. **D)** Subcellular localization of mCerulean-PR. mCerulean-PR localizes to the nucleus, with a preference for the nucleolus (top panels), but when Kap114 is co-expressed mCerulean-PR becomes more diffusely nuclear. **E)** Frequency of nucleolar (D top panel) or nuclear diffuse (D bottom panel) mCerulean-PR signal observed in cells expressing mCerulean-PR or co-expressing mCerulean-PR and Kap114. Total numbers of cells counted indicated.

When comparing the net charge of all yeast NTRs by subtracting the positively charged residues (Arg/Lys) from the negatively charged residues (Asn/Glu), it is striking that Kap114 (in orange in Fig 7C) and Sxm1, and Kap123 from the transport cluster 3, are similarly strongly negatively charged with net negative charges of 65, 59 and 75, respectively. The exportins are on the left side of the graph, showing RNA exportins Mex67-Mtr2 with net positive charges, and Cse1, Crm1, and Los1 with a net negative charge of 28 and 29, respectively (in magenta in Fig 7C). Kap120, from cluster 4 is present on the left side of this graph as well with a net charge of -32 (in orange in Fig 7C).

The direct binding of polyPR and Kap114 predicted by the simulations made us wonder why Kap114-GFP and polyPR were not observed to colocalize in cells (Fig S1B). We tested the possibility that the normal cellular levels of Kap114 may be too low to support stable association with polyPR. We thus overexpressed Kap114 and followed the subcellular localization of mCerulean-PR. We indeed see that overexpression of Kap114 relocalized mCerulean-PR from the nucleolus. Specifically, without overexpression of Kap114, mCerulean-PR is nucleolar in 82% of cells, but with overexpression of Kap114, mCerulean-PR has a nuclear diffuse localization in 77% of cells (Fig 7D/E).

We conclude that the distinct transport profile of Kap120Δ cells compared to the transport profiles of the other NTR knockouts and cells expressing polyPR (Fig 6), and the prediction of ionic interaction between polyPR and negatively charged NTRs (Fig 7), can be taken as an indication that the mechanism through which polyPR derails NCT is by depleting the cells of functional NTRs.

## Discussion

In this study we comprehensively analyze the transport defects arising from expression of C9ALS associated DPRs and compare them to those occurring in several stress conditions and mutants. The systematic quantitative assessment of 100 different transport readouts from four transport reporters revealed, first of all, that DPR expression impacts transport about as much as exposure to common environmental stresses. Projecting the data from our four transport receptors to the whole proteome, our data predicts a chronic global mislocalization of many proteins, which could have severe effects on cellular viability. A comparison of the specific patterns of alterations in transport related to the expression of polyPR and polyGA, to those obtained when cells were exposed to different stress situations, revealed striking similarities. The transport phenotype of polyPR expressing cells clusters with those measured in four NTR deletion mutants, while the transport phenotype of polyGA expressing cells clusters with stresses that affect energy maintenance. If we consider that changes in NCT may serve as a fingerprint reporting on cell physiology, then we may infer that the cellular mechanisms compromised by polyPR or polyGA expression are related to the maintenance of, respectively, NTR availability and energy levels.

Our study thoroughly characterizes the yeast model expressing PR_50_ or GA_50_, in terms of toxicity and changes of the NCT machinery. It recapitulates the toxicity and localization of the DPRs as reported in other models, where PR_50_ is toxic and localized in the nucleolus and GA_50_ is often not toxic and forms aggregates in the cytosol^35–39,48,59,60,63–65^ (and reviewed in^40^). Our analysis of the abundance and localization of the components of the nuclear transport machinery, including NPC components, almost all NTRs, and the components of the Ran gradient, shows that expression of polyGA and polyPR does not induce significant changes. Our data does not support blocking of NPCs, nor does it show prominent colocalization of DPRs with NTRs or the NPC as previously reported^52,87,88^. At the same time, we do find that the expression of both DPRs impact the transport by different NTRs (Crm1, Kap60/Kap95, Kap104, and Kap121/Pse1) to varying degrees and directions. Specifically, polyPR increases the transport by Kap60/95, and Crm1, but decreases transport of the other two import routes. In contrast, polyGA reduces transport by Kap60/95 and Crm1, but does not impact Pse1 transport.

We found that the transport phenotype of polyPR expressing cells clusters with those measured in four NTR deletion mutants. We propose that the similarity between the NCT profiles measured upon the removal of one NTR -as in the knockouts- and the polyPR expressing strains may mechanistically be explained by the electrostatic interaction of polyPR with NTRs, as shown in our coarse-grained molecular dynamics simulations. Such direct biding of polyPR to NTRs is in line with previous reports^51,55^. The binding of polyPR to NTRs may leave these NTRs unsuited to bind their cargo and/or the FG-Nups of the NPC, thereby compromising transport. Following this logic, both polyPR expression and deletion of the more negatively charged NTRs, Sxm1, Kap122, and Kap123, result in a reduction of the pool of functional NTRs, and thus in similar transport defects.

The similarity in NCT phenotypes of polyGA expressing cells and diverse environmental challenges (glucose deprivation, oxidative stress, and heat stress) to which the cells have evolved adaptive mechanisms might explain why polyGA expression is not toxic. It may also suggest that the transport defects arise secondary to the stress associated with polyGA expression. It would be interesting to compare these transport phenotypes to those obtained with the expression of other non-toxic aggregation prone proteins; if similar this may reflect the profile of a cell in which protein homeostasis is challenged.

How do we value our data towards the understanding of C9ALS? Specifically related to the multifactorial nature of C9ALS, where the G_4_C_2_ sequence of the genome, the RNAs that are transcribed as well as the 5 different DPRs all exert their effects, a direct translation for the reductionists approach taken here is not possible. In addition, the yeast model is obviously very distant from humans. However, due to the conservation of basic biology, including the NCT system, we can infer that our data suggest that also in patients there will be a global impact on the kinetics of transport. Quantitatively, the impact on transport is expected to be in the same order as those measured in different physiologically relevant stresses including ageing. The quantitatively modest effects on NCT may be the explanation for the diverse outcomes of transport studies in the literature (reviewed in^40^). As to the question of causality, our data supports that some aspects of the C9orf72 pathology are indirect while other are related to direct interactions with the NCT machinery. Whether such direct interactions with the NCT provide opportunities for future interventions should be investigated more thoroughly and preferably these studies should go hand-in-hand with studies investigating the robustness of NCT under stress and ageing.

## Materials and methods

### Strains and growth conditions

All *Saccharomyces cerevisiae* strains used in this study have the BY4741 genotype, and are listed in Table 4. For the fluorophore-tagging of DPRs mCerulean3 was PCR amplified from CrGE2 (Crowding sensor^89^) and inserted between GAL1 promotor and PR_50_ sequence via BamHI/PstI sites. The ATP sensor was constructed by integrating a synthetic pTEF1-his6-ymEGFPΔ11-B.subtilis ε construct (GeneArt, ThermoFischer, as based on^70^) into a pRS303-ymScarletI vector using SpeI/NcoI sites. Apart from the strains in Fig 1C/D, all DPR containing strains were transformed with pRS416-PR_50_/GA_50_. Fig 1C/D was made with the mCerulean-tagged DPRs. GFP-tagged strains were taken from the 4000-GFP yeast library (Thermofisher), while knockout strains were taken from the Yeast Knockout Collection (Invitrogen).

For expression of DPRs and transport reporter proteins, cells were grown at 30°C, with shaking at 200 RPM using the appropriate Synthetic Dropout medium supplemented with 2% glucose in overnight culture. The next day cultures were diluted 1:10 in SD medium supplemented with 2% D-raffinose, and again for an overnight culture. Expression of DPRs and transport reporter proteins was induced with 0.1% galactose 3h prior to the start of an experiment, while the cultures had an OD_600nm_ between 0.6-0.9. For the different stress conditions the transport reporters were induced with 0.1% galactose for 1h.

### Spot assay

Cells were grown overnight in 2% glucose containing medium, diluted the next day 1:10 in 3ml 2% raffinose medium. At the end of the day cells were diluted to obtain a culture of OD_600nm_ ∼0.3 the following morning. This culture was used to dilute cells in sterilized water to obtain 10^4^–10^6^ cell/ml densities, and then 5μl was spotted on the appropriate YPD or YPGal plates.

### Microscopy

Microscopy was performed at 30°C on a Delta Vision Deconvolution Microscope (Applied Precision), using InsightSSITM Solid State Illumination of 435 and 525 nm, an Olympus UPLS Apo 40x or 100x oil objective with 1.4NA and softWoRx software (GE lifesciences). Detection was done with a CoolSNAP HQ2 or PCO-edge sCMOS camera.

### Microfluidic devices

To determine division times for the PR_50_/GA_50_ expressing strains, an ALCATRAS chip was used as previously detailed^62^. DIC images of cells were taken every 20 min. to follow all divisions of each cell. The efflux assay was performed in the CellASIC Onix microfluidic perfusion system (Merck Millipore). Cells were loaded 3 times 5 sec. at 8psi or 5psi for smaller cells, medium was run at 2psi to flush out free cells. Medium was exchanged for the appropriate synthetic dropout medium supplemented with 10 mM sodium azide and 10 mM 2-deoxy-D-glucose^90^, additionally ponceau S was used to allow fluorescence in the A594 channel to determine when the poison front hit the cells. The addition of sodium azide and 2-deoxyglucose depletes the cell of energy and destroys the Ran-GTP/GDP gradient thus abolishing active transport of reporter proteins. We measured the net efflux of reporter proteins by imaging the cells every 2,5 millisecond for a period of 20 seconds in the FITC channel starting 5 seconds after the poison front.

### Data analysis of N/C ratios and efflux rates

Microscopy data was quantified with open source software Fiji (https://imagej.net/Welcome;^91^). Fluorescent intensity measurements were corrected for background fluorescence. To quantify the nuclear localization (N/C ratio) of the reporters, the average fluorescence intensity at the nucleus and cytosol were measured. The nuclear area was determined using live Hoechst33342 stain (NucBlue Live Cell Stain Ready Probes, Invitrogen) or using the signal of the GFP-reporters in case of the poison assay. A section in the cytosol devoid of vacuoles was selected for determining the average fluorescence intensity in the cytosol. Leak rates were determined by linear regression lines in GraphPad Prism (version 8.3.0). Cells with linear regression lines with standard deviation of the residuals (Sy.x) above 0.03 were excluded from the analysis. Statistical parameters including the definitions and exact values of N, distributions and deviations are reported in the figures and corresponding figure legends. Each single cell counts as a biological replicate, and grey colours in the figures 3,4, and 5 are different cultures. Significance of changes were determined with a Mann-Whitney test, since not all measured conditions were normally distributed.

### Stress interventions

All stress interventions were performed after induction of the transport reporters with 0.1% galactose for 1h, and additionally incubated for 5 minutes with NucBlue (NucBlue Live Cell Stain Ready Probes, Invitrogen), to allow for determination of the nuclear area.

Heat shock was performed after growing the cells as previously stated, then harvesting 1.5ml cells and replacing the medium with preheated medium at the appropriate (37°C/42°C/46°C) temperature. Next the culture was maintained at this temperature for 5 or 10 minutes (as indicated) in a water bath, after which cells were immediately imaged.

Starvation was performed either by growing the culture to saturation in medium with 2% raffinose over 20 hours, after which the reporters were induced with 0.1% galactose for 1h. For the water starvation experiment, we followed the method as previously described^92^. In short, GFP-reporter strains were grown in SD medium supplemented with 2% glucose and grown overnight. The next day, cells were diluted in Sgal medium with 4% D-galactose and grown to an OD_600nm_ between 0.6 and 0.9. Then cells were harvested and resuspended in water, and maintained shaking at 30°C. Samples were taken from the galactose culture at t=0, and from the water culture after 30min, followed by a sample after 24hr.

Osmotic shock was performed as described previously^82^. In short, 1.5ml of cells was harvested after induction of the transport reporters, and resuspending in either 100μl low osmolality buffer (50mM NaPi, pH 7), or high osmolality buffer (50mM NaPi, 1M NaCl) to induce osmotic upshift.

Oxidative stress was performed by adding 4mM H_2_O_2_ (Sigma-Aldrich) to 1.5ml of induced cells and imaging directly, or by adding 1.2mM menadione (Sigma-Aldrich) for 40 minutes overlapping with the galactose inducing.

15% (v/v) ethanol was added during the last 20minutes of galactose induction.

### Cluster analysis of interventions

For each intervention, the median fold change compared to wild type, on log2 scale, was used in a Principal Component Analysis (prcomp, centered and scaled; R studio version 1.4.1717). This data was used for the standard function of the biplot and the heatmap of R. The clustering was performed with the data.kmeans function (5 clusters, iter.max = 10, n = 25).

### Proteomics sample preparation

10ml cultures of wild type, PR_50_- and GA_50_-repeat expressing cells were grown as described. Cells were harvested, washed twice in PBS, and lysated by shaking with glass beads according to the yeast protocol in the FastPrep machine (MpBio). Yeast lysate samples were mixed with LDS loading buffer (NuPAGE) at a concentration of 10 ug total protein in a total volume of 20 μL. The sample was run briefly into a precast 4-12% Bis-Tris gels (Westburg, ran for maximally 5 min at 100 V). The band containing all proteins was visualized with Biosafe Coomassie G-250 stain (Biorad) and excised from gel. Small pieces were washed subsequently with 30% and 50% v/v acetonitrile in 100 mM ammonium bicarbonate (dissolved in milliQ-H_2_O), each incubated at RT for 30 min while mixing (500 rpm) and lastly with 100% acetonitrile for 5 min, before drying the gel pieces at 37 °C. The proteins were reduced with 30 μL 10 mM dithiothreitol (in 100 mM ammonium bicarbonate dissolved in milliQ-H_2_O, 30 min, 55 °C) and alkylated with 30 μL 55 mM iodoacetamide (in 100 mM ammonium bicarbonate dissolved in milliQ-H_2_0, 30 min, in the dark at RT). The gel pieces were washed with 100% acetonitrile for 30 min while mixing (500 rpm) and dried in at 37 °C before overnight digestion with 30 μL trypsin (1:100 g/g, sequencing grade modified trypsin V5111, Promega) at 37 °C. The next day, the peptides were eluted from the gel pieces with 30 μL 75% v/v acetonitrile plus 5% v/v formic acid (incubation 20 min at RT, mixing 500 rpm). The elution fraction was diluted with 900 μL 0.1% v/v formic acid for cleanup with a C18-SPE column (SPE C18-Aq 50 mg/1mL, Gracepure). This column was conditioned with 2×1 mL acetonitrile plus 0.1% v/v formic acid, and re-equilibrated with 2×1 mL 0.1% v/v formic acid before application of the samples. The bound peptides were washed with 2×1mL 0.1% v/v formic acid and eluted with 2×0.4 mL 50% v/v acetonitrile plus 0.1% v/v formic acid. The eluted fractions were dried under vacuum and resuspended in 0.1% v/v formic acid to a final concentration of around 1 μg/μL total protein starting material.

### Targeted proteomics analyses

Selected reaction monitoring (SRM) analyses were performed on a triple quadrupole mass spectrometer with a nano-electrospray ion source (TSQ Vantage, Thermo Scientific) for the peptides listed in Table 2, for which isotopically labeled peptides were derived from generated QconCATs (these QconCATs were kindly provided by Prof. R. Beynon, University of Liverpool UK). Chromatographic separation of the peptides was performed by liquid chromatography on a nano-UHPLC system (Ultimate UHPLC focused, Thermo Scientific) using a nano column Acclaim PepMapC100 C18, 75 μm x 50 cm, 2 μm, 100 Å, Thermo Scientific). Samples were injected at a concentration of 1 μg protein and an amount of isotopically-labeled standards optimized for the chosen application using the μL-pickup system using 0.1% v/v formic acid as a transport liquid from a cooled autosampler (5 °C) and loaded onto a trap column (μPrecolumn cartridge, Acclaim PepMap100 C18, 5 μm, 100 Å, 300 μm id, 5 mm Thermo Scientific). Peptides were separated on the nano-LC column using a linear gradient from 3-60 % v/v acetonitrile plus 0.1% v/v formic acid in 100 minutes at a flowrate of 300 nL/min. The mass spectrometer was operated in the positive mode at a spray voltage of 1500 V, a capillary temperature of 270 °C, a half maximum peak width of 0.7 for Q1 and Q3, a collision gas pressure of 1.2 mTorr and a cycle time of 1.2 ms. Optimal collision energies (CE) were predicted using the following linear regressions: CE = 0.03*m/z precursor ion + 2.905 for doubly charged precursor ions, and CE=0.03*m/z precursor ion +2.467 for triply charged precursor ions. For each of the peptides, the optimal precursor charge and three optimal transitions were selected after screening with the QconCAT peptides. The measurements were scheduled in windows of 5 minutes around the pre-determined retention time. For NTR, Nup, and Ran cycle component abundances were calculated as amount compared to total protein amounts. Then the average of three runs, one being a technical and one a biological replicate, each in triplicate, was used to normalize PR- and GA-expressing cell abundances to WT amounts (Source data 1, for biological and technical replicates).

**Table 2.**
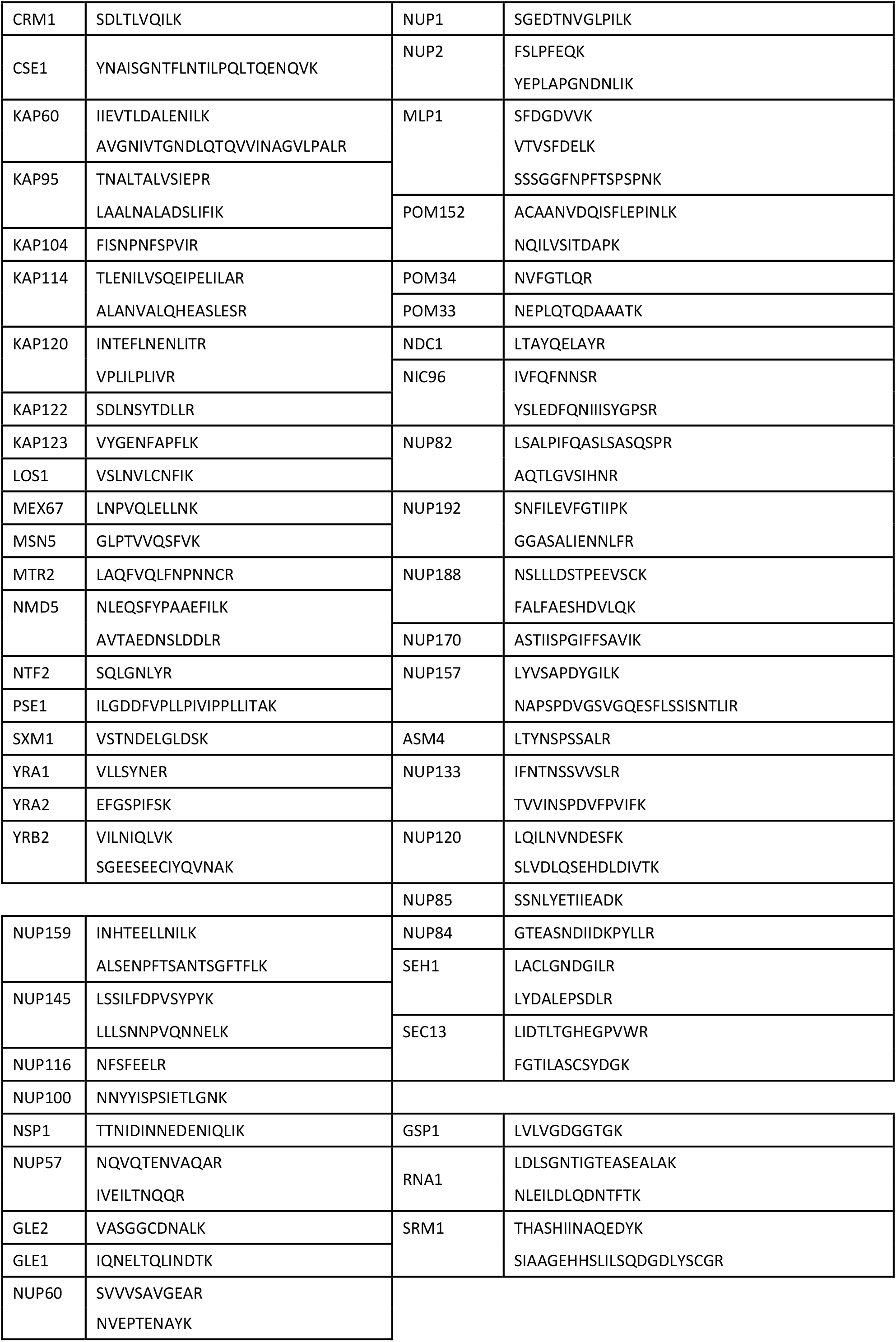
SRM mass spectrometry peptides in this study.

### Discovery-based proteomics analyses

Discovery mass spectrometric analyses were performed on an orbitrap mass spectrometer with a nano-electrospray source (Orbitrap Q Exactive Plus, Thermo Scientific). Chromatographic separation of the peptides was performed by liquid chromatography on a nano-HPLC system (Ultimate 3000, Thermo Scientific) using a nano-LC column (Acclaim PepMapC100 C18, 75 μm x 50 cm, 2 μm, 100 Å, Thermo Scientific). In general, an equivalent of 1 μg total protein starting material was injected using the μL-pickup method with 0.1% v/v formic acid as a transport liquid from a cooled autosampler (5 °C) and loaded onto a trap column (μPrecolumn cartridge, Acclaim PepMap100 C18, 5 μm, 100 Å, 300 μmx5 mm, Thermo Scientific). Peptides were separated on the nano-LC column using a linear gradient from 2-45% v/v acetonitrile plus 0.1% v/v formic acid in 85 min at a flowrate of 300 nL/min. The mass spectrometer was operated in the positive mode in a data-dependent manner, with automatic switching between MS and MS/MS scan using a top-15 method. MS spectra were acquired at a resolution of 70.000 with a AGC target of 3e^6^ ions or a maximum integration time of 50 ms at a scan range of 300 to 1650 m/z. Peptide fragmentation was performed with higher energy collision dissociation (HCD) with the energy set to 28 NCE. The intensity threshold for ions selection was set at 2.0e^4^ with a charge exclusion of 1≤ and ≥6. The MS/MS spectra were acquired at a resolution of 17.500, with a target value of 1e^5^ ions or maximum integration time of 50 ms and the dynamic exclusion set to 20 sec.

LC-MS raw data were processed with MaxQuant (version 1.5.2.8^93^). Peptide and protein identification was carried out with Andromeda against a yeast SwissProt database (www.uniprot.org, 6721 entries, downloaded on June 2020). For peptide identification two miss cleavages were allowed, a carbamidomethylation on cysteine residues as a static modification and an oxidation of methionine residues and acetylation of protein N-termini as variable modifications. Peptides and proteins were identified with an FDR of 1%. For a protein identification at least one unique peptide had to be detected and the match between runs option was enabled. Proteins were quantified with the MaxLFQ algorithm^94^ considering only unique peptides and a minimum ratio count of one. Results were exported as tab-separated *.txt for further data analysis.

For cargo abundances, only one exploratory MS run in triplicate was performed, therefore only LFQ intensities were averaged over one set of triplicates, and compared between WT, PR-, and GA-expressing cells (Source data 3).

### Coarse-grained model

We use a modified version of the implicit solvent, coarse-grained one-bead-per-amino acid (1BPA) model developed earlier^95^. The coarse-grained models of Kap120 (PDB:6fvb) and Kap114 (PDB:6aho) are built by considering beads at the position of α-carbons in the crystal structures and introducing a network of stiff harmonic bonds that maintains the secondary and tertiary structure of the NTRs. This network of bonds is represented by the harmonic potential *ϕ*_network_ = *K*(*r* − *b*)^2^, where *K* is 8000 kJ/nm2 and *b* is the original distance between the amino acid beads in the crystal structure. A bond is made between the beads if *b* is less than 1.4 nm. The missing regions in the crystal structures of Kap120 and Kap114 have around 75% of their residues in a coil conformation calculated using PSIPRED^96^, and are included in the coarse-grained model as disordered regions.

For polyPR-polyPR interaction, we take into account hydrophobic/hydrophilic and electrostatic interactions. The hydrophobic/hydrophilic interactions are represented by:

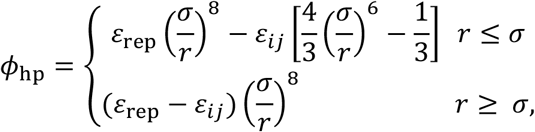

where 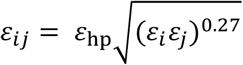 is the strength of the interaction for each pair of amino acids (*i,j*) and *σ* = 0.6 nm. The values of *ε*_hp_ and *ε*_rep_ are 13 and 10 kJ/mol, respectively. The relative hydrophobic strength values (*ε*_*i*_ ∈ [0,1]) of the different amino acids are listed in Table 3. The hydrophobic strength values of charged residues are slightly increased compared to the original model^95^ in line with our recent work^86^.

**Table 3.** Relative hydrophobic strength values of the different amino acids

| Amino acid | $\epsilon_i$ | Amino acid | $\epsilon_i$ |
| --- | --- | --- | --- |
| A | 0.7 | L | 1 |
| R | 0.005 | K | 0.005 |
| N | 0.33 | M | 0.78 |
| D | 0.005 | F | 1 |
| C | 0.68 | P | 0.65 |
| Q | 0.64 | S | 0.45 |
| E | 0.005 | T | 0.51 |
| G | 0.41 | W | 0.96 |
| H | 0.53 | Y | 0.82 |
| I | 0.98 | V | 0.94 |

**Table 4.**
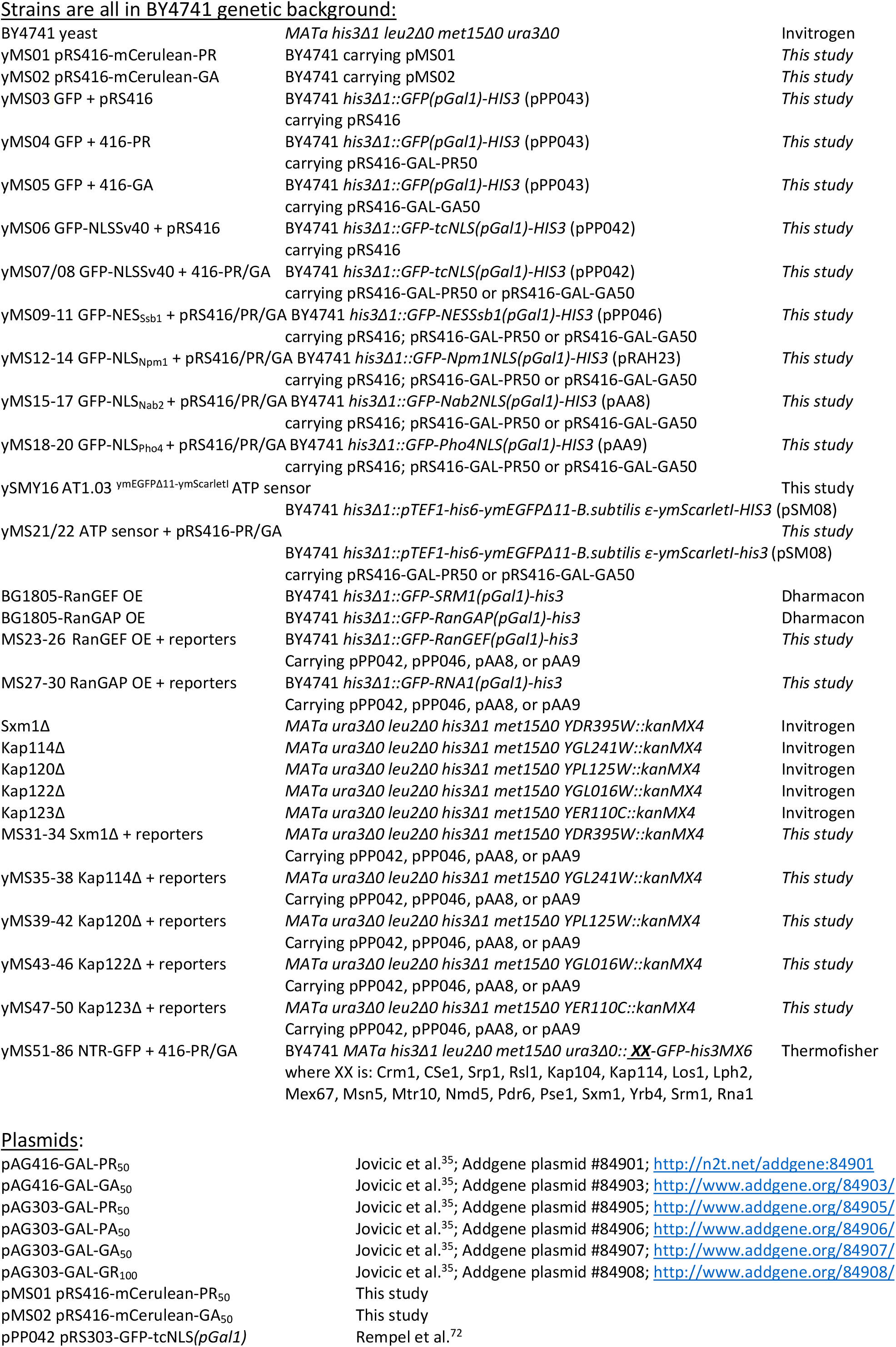
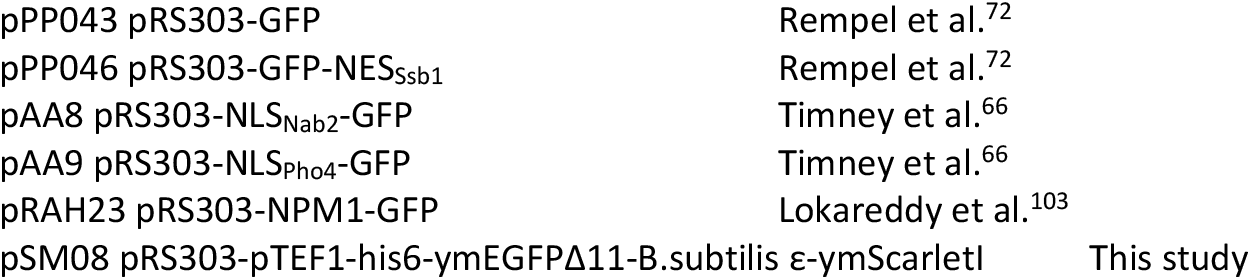
Yeast strains used in this study.

The electrostatic interactions between charged residues are described by the modified Coulomb law:

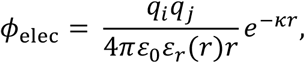

where 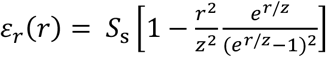 is the distance-dependent dielectric constant of the solvent with *S*_*s*_ = 80 and *z* = 0.25 nm. The value of the Debye screening coefficient, *k*, is 1 nm^-1^ for monovalent salt concentration *C*_salt_ = 100 mM, and 1.5 nm^-1^ for *C*_salt_ = 200 mM.

The interactions between polyPR and NTRs can be classified in three categories: electrostatic interactions, cation-pi interactions, and excluded volume interactions. For electrostatic interactions, we use the same electrostatic potential (*ϕ*_elec_) as described above. To take into account the cation-pi interactions between Arginine (in polyPR) and the aromatic residues F,Y, and W (in the NTRs), we use an 8-6 Lennard-Jones (LJ) potential that replaces *ϕ*_hp_ for the RF, RY, and RW interactions:

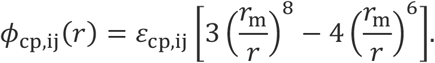

The parameter *r*_m_, which is the distance at which *ϕ*_cp,ij_ reaches its minimum value, is set to 0.45 nm equal to the weighted average distance between the guanidinium group of Arginine and an aromatic ring at different orientations (Planar, Oblique, Orthogonal)^97^. To find *ε*_cp,ij_ for the different cation-pi pairs, we set the RY cation-pi energy as a basis for calculating the cation-pi energies for RF and RW using PDB statistics. According to all-atom free energy calculations, the RY interaction energy is comparable to the strongest interaction between different non-charged residues at physiological salt concentrations^98^. Therefore, in order for the cation-pi interactions to be compatible with the 1BPA force field, we set *ε*_cp,RY_ = 5 kJ/mol which is similar to the deepest potential depth in the 1BPA force field (5.2 kJ/mol). To estimate the energy difference between RY and the other combinations, similar to^99^, we use the PDB cation-pi contact frequencies in an aqueous environment^100^, and a formulation for statistical potential^99,101^. In a dataset analyzed in^100^, the frequencies of R, F, Y, and W are *p*(R) = 10919, *p*(F) = 9162, *p*(Y) = 8309, *p*(W) = 3412, respectively, and the cation-pi contact frequencies for RF, RY, and RW are *p*(RF) = 630, *p*(RY) = 749, *p*(RW) = 609. Based on these values the energy differences between different cation-pi pairs can be estimated. As an example, using *k*_B_*T* ≈ 2.5 kJ/mol at *T* = 300 K, the energy difference between RY and RF (former minus latter) can be estimated as −*k*_B_*T*ln([*p*(RY)/*p*(RF)][*p*(F)/*p*(Y)]) ≈ −0.7 kJ/mol. The value of *ε*_cp,RF_ is then *ε*_cp,RF_ ≈ *ε*_cp,RY_ − 0.7 kJ/mol ≈ 4.3 kJ/mol. A similar calculation results in *ε*_cp,RW_ = 6.70 kJ/mol.

For the hydrophilic/hydrophobic interactions between poly-PR and the rest of the NTR residues (the orange residues in Fig 7A), we use *ϕ*_hp_ with *ε*_ij_ = 10 kJ/mol which leads to an excluded volume potential that vanishes at *r* = 0.6 nm. For the interactions between the residues within the disordered regions of NTRs, we use the 1BPA force field featuring *ϕ*_hp_ and *ϕ*_elec_ as described above.

### Simulation and contact analysis

Langevin dynamics simulations are performed at 300 K at monovalent salt concentrations of 100 mM and 200 mM in NVT ensembles with a time-step of 0.02 ps and a Langevin friction coefficient of 0.02 ps^-1^ using GROMACS version 2018. Simulations are performed for at least 2.5 μs in cubic periodic boxes, and the last 2 μs are used for the contact analysis. The time-averaged number of contacts *C*_*t*_ between the polyPR and NTRs is obtained by summing the number of contacts per time frame (i.e. the number of polyPR/NTR residue pairs that are within 1 nm) over all frames and dividing by the total number of frames. The error bars in Fig 7B are half of the standard deviation. The structures in Fig 7 are drawn using VMD^102^.

## Abbreviations

C9ALS: Chromosome 9 Amyotrophic Lateral Sclerosis
DPRs: DiPeptide Repeat proteins
FG-Nup: phenylalanine–glycine repeat containing Nup
FUS: Fused in Sarcoma
GA/GP/GR/PA/PR: Glycine-Alanine, Glycine-Proline, Glycine-Arginine, Proline-Alanine, Proline-Arginine
NCT: NucleoCytoplasmic Transport
NES: Nuclear Export Signal
NLS: Nuclear Localization Signal
NPC: Nuclear Pore Complex
NTF: Nuclear Transport Factor
NTR: Nuclear Transport Receptors
Nup: Nucleoporin
RAN: translation Repeat-Associated Non-AUG translation
ROS: Reactive Oxygen Species
TDP43: TAR DNA-binding protein-43

## Acknowledgments

This work was financially supported by the Netherlands Organization for Scientific Research (NWO BBOL 737.016.016 to LMV and P.R.O) and NWO-*Vici* (VI.C.192.031) to LMV. We thank Thamar Jessurun Lobo and Prof. Victor Guryev for their help with the PCA and cluster analysis. We thank Prof. Michael Chang and all members from the Veenhoff and Chang laboratories for their critical input in the project.

## Source data files

1. SRM proteomics data, figure 2
2. Untargeted proteomics data, cargo abundances
3. Raw N/C ratios, figure 3
4. Raw N/C ratios, figure 4
5. Raw N/C ratios, figure 5
6. NTR charge, figure 7

## Supplementary

**Fig 2 – figure supplement 1.**
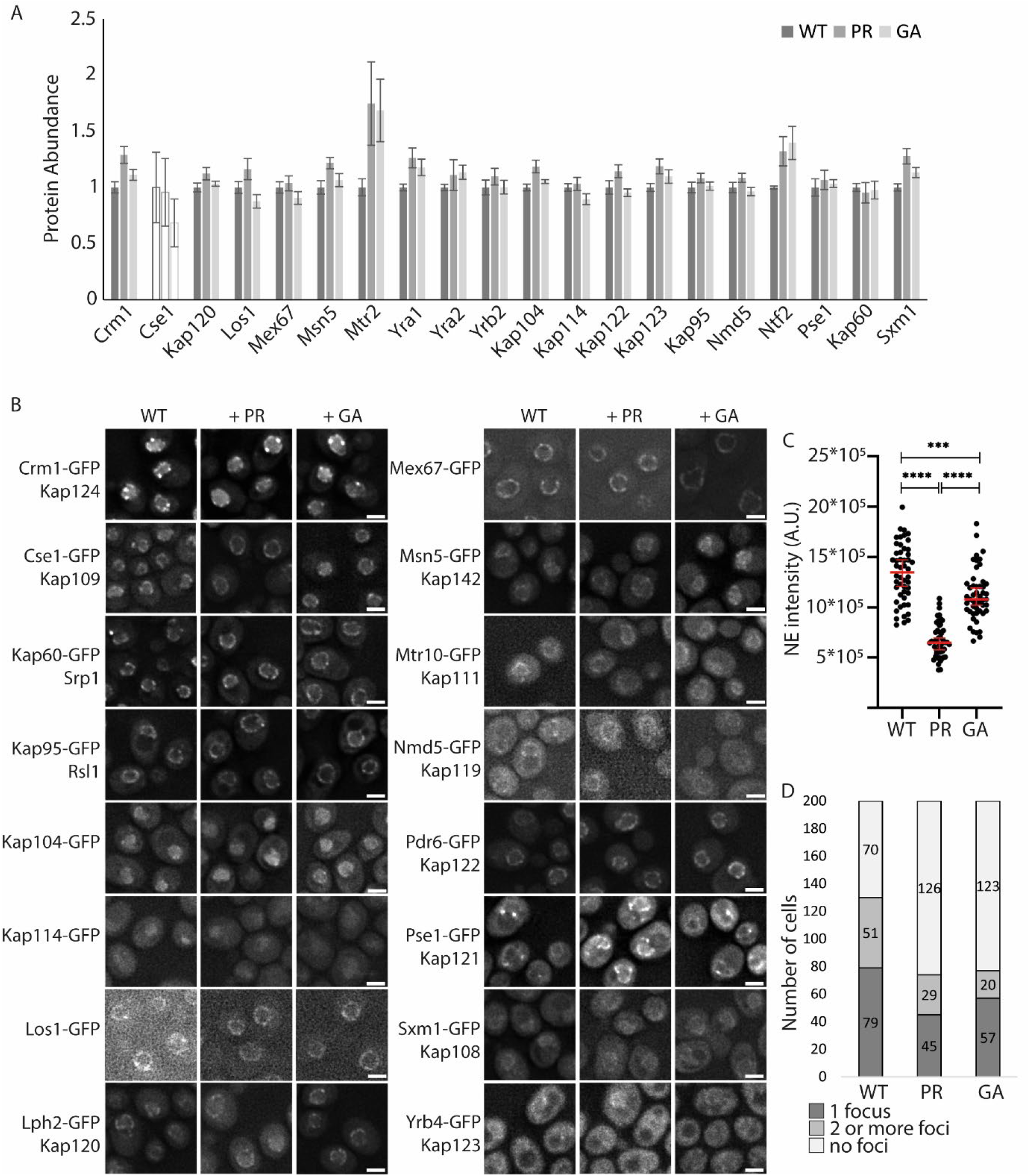
Abundance and localization of NTRs in polyPR and polyGA expressing cells. **A)** Abundance of NTRs in whole cell extracts of WT or PR_50_/GA_50_-expressing cells determined by SRM-based proteomics in two biological and one technical replicate, except for Cse1 which has one replicate (in white bars). **B)** Localization of GFP-tagged NTRs expressed from native promotor and genomic location compared between WT cells and cells expressing PR_50_ or GA_50_; scale bar equals 2μm. **C)** The intensity of Cse1-GFP at the nuclear envelope in WT, PR-expressing, and GA-expressing cells (all 50 cells). **D)** The number of Pse1-GFP foci in WT cells, or those expressing PR/GA.

**Fig 2 – figure supplement 2.**
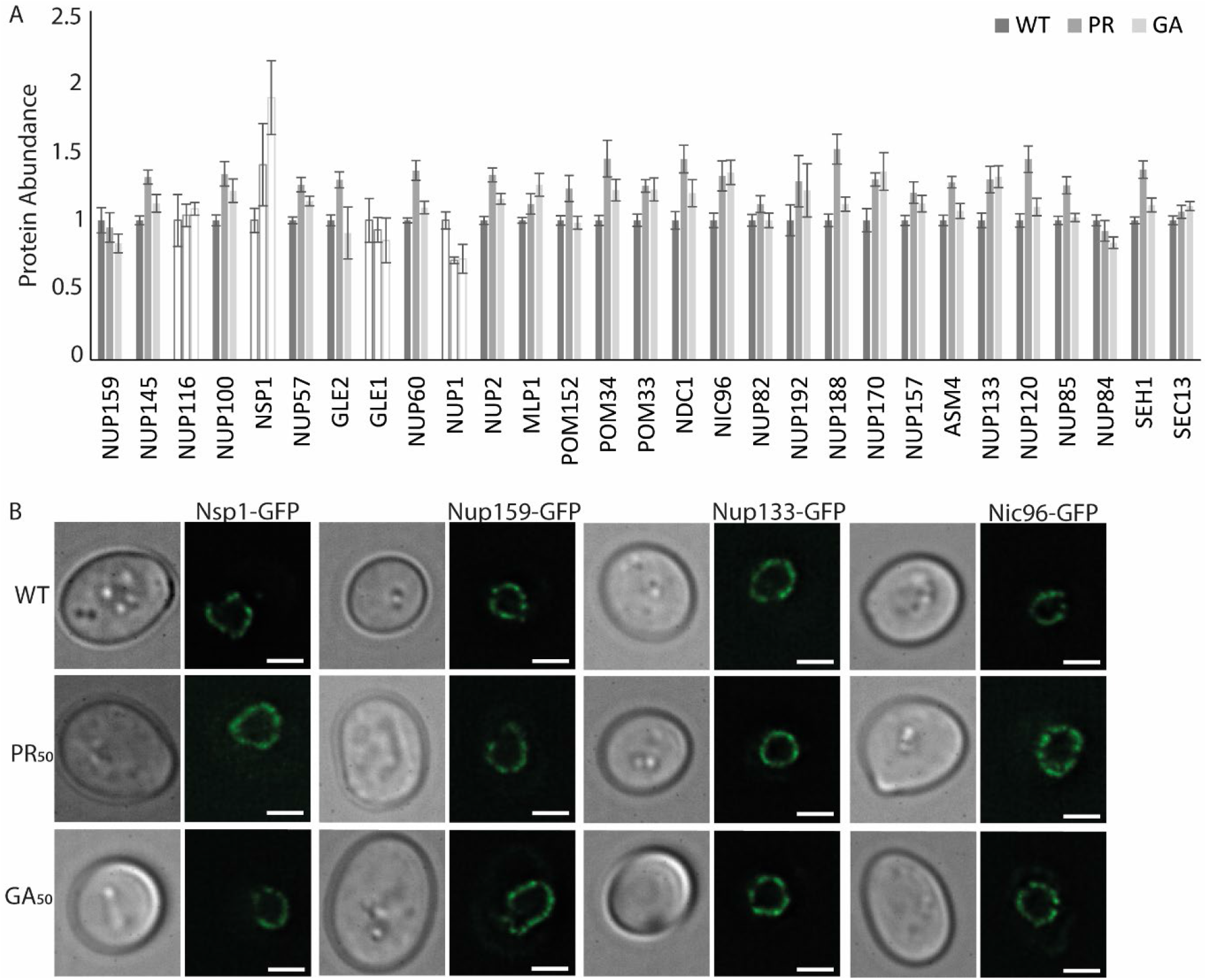
Abundance and localization of nucleoporins in polyPR and polyGA expressing cells. **A)** Abundance of nucleoporins in whole cell extracts of WT or PR_50_/GA_50_-expressing cells determined by SRM-based proteomics in two biological and one technical replicate, except for Nup116, Nsp1, Gle, Nup1 which have one replicate (in white bars). **B)** The localization of GFP-tagged Nsp1, Nup133, Nup159, and Nic96 does not change upon PR_50_ expression; Nic96 and Nup133 also shown in Fig2. Scale bar equals 2μm.

**Fig 2 – figure supplement 3.**
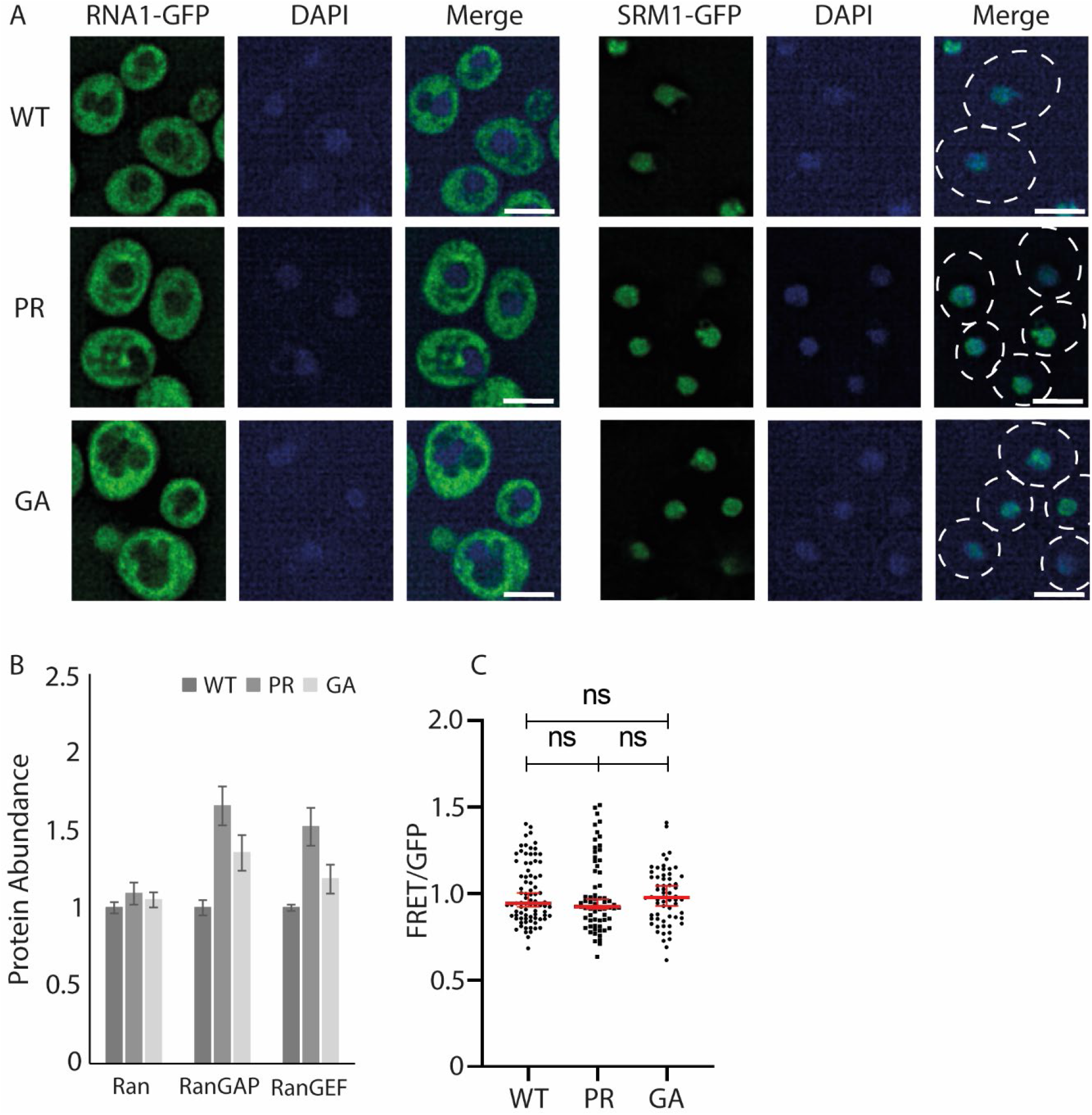
Abundance and localization of Ran, RanGAP and RanGEF in polyPR and polyGA expressing cells. **A)** The localization of GFP-tagged RanGAP and RanGEF does not change with PR_50_ expression. RanGAP is cytoplasmic diffuse and RanGEF nuclear diffuse. Nuclear staining with DAPI, scale bar equals 2μm. Also shown in Fig 2. **B)** Abundance of Ran, RanGAP and RanGEF in whole cell extracts of WT or PR_50_/GA_50_-expressing cells determined by SRM-based proteomics in two biological and one technical replicate. **C)** A FRET-based ATP-sensor measures free ATP levels. The FRET over GFP ratio measured in live cells is not significantly changed between WT (n = 83) and PR-expressing cells (n = 67) or GA-expressing cells (n=62) as determined via the Mann-Whitney test.

**Fig S1.**
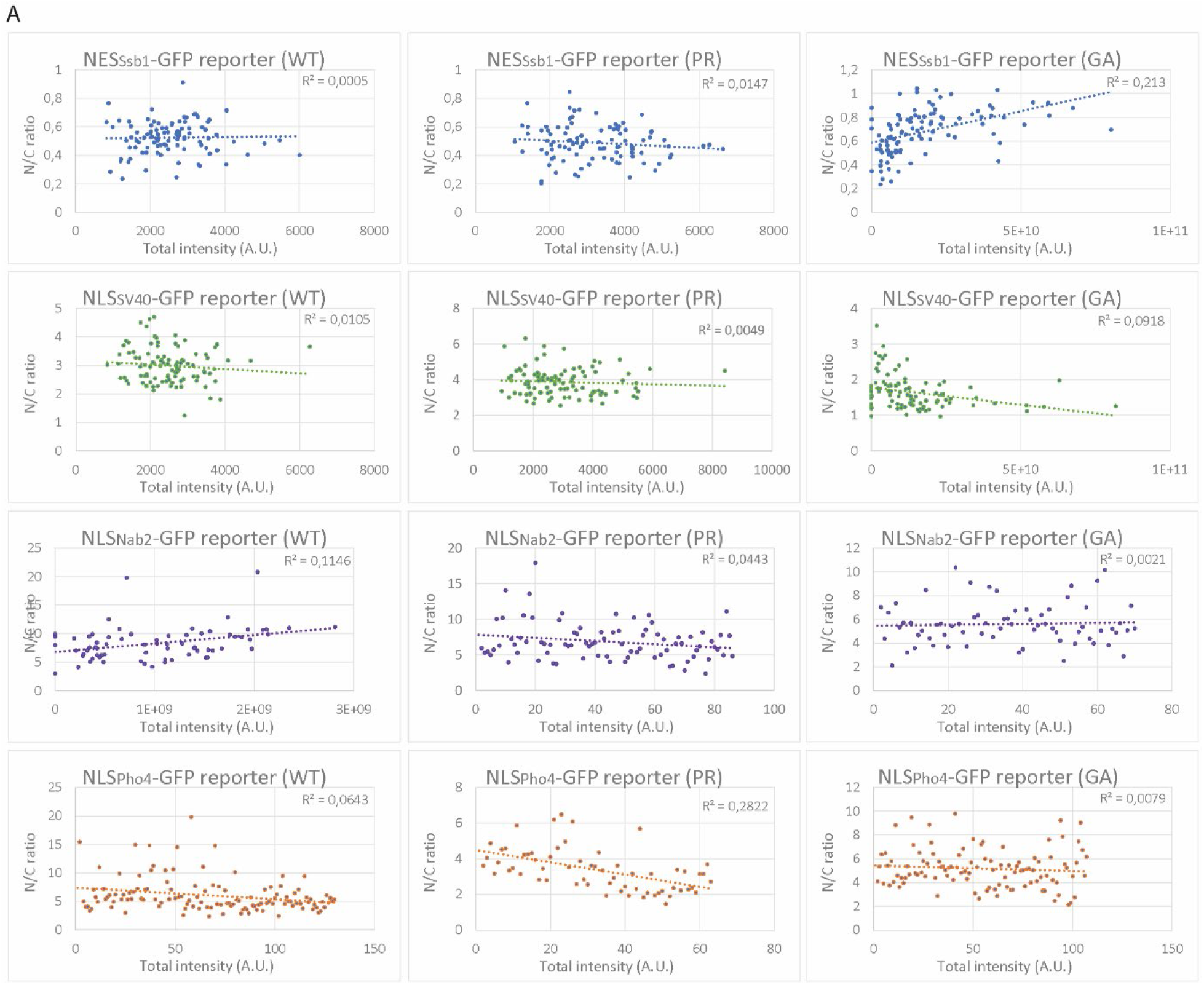
Nuclear accumulation of transport reporters is independent of their expression level. **A)** N/C ratios from WT, PR_50_- and GA_50_-expressing cells show no correlation between expression level (total intensity of both N and C) and N/C ratios.

**Table S1.** Characteristics of NTRs studied

| **Table S1** Characteristics of NTRs studied |  |  |  |  |  |
| --- | --- | --- | --- | --- | --- |
| NLS | Ssb1 NES | Sv40 NLS | Npm1 NLS | Nab2 NLS | Pho4 NLS |
| Karyopherin | Crm1 | Kap60/95 | Kap60/95 | Kap104 | PSE1 |
| Human NTR homologue | Exportin | 60: Karyopherin $\alpha$<br>95: Karyopherin $\beta$ | 60: Karyopherin $\alpha$<br>95: Karyopherin $\beta$ | TNPO1 | IPO5 |
| Localization NTR (see Fig 2 – figure supplement 1B) | Nuclear + foci at nuclear ring | Nuclear ring | Nuclear ring | Nuclear | Foci at nuclear ring |
| Abundance NTR (molecules/cell) (according to SGD yeastgenome.org) | 8276 ± 2396 | 60: 8820 ± 4085<br>95: 15792 ± 5198 | 60: 8820 ± 4085<br>95: 15792 ± 5198 | 4755 ± 2218 | 8125 ± 1774 |
| Used localization signal sequence | IEAALSDALA<br>ALQI<sup>75</sup> | PKKKRKVDPKK<br>KRKV<sup>76</sup> | AVKRPAATKK<br>AGQKKKKLD<sup>77</sup> | DNSQRFTQRG<br>GGAVGKNRRG<br>GRGGNRGGRN<br>NNSTRFNPLAK<br>ALGMAGESN<sup>78</sup> | ANKVTKNKSN<br>SSPYLNKRRGK<br>PGPDSATSLFE<br>LPDSVIPTPKP<br>KPKPKNTPKVILP<sup>79</sup> |
| Binding affinity to NLS | X | 60: 10nM<sup>104</sup><br>Complex: 35nM<sup>105</sup><br>Impα to Impβ: 11-14nM<sup>105</sup> | 60: 46 ± 14nM<sup>103</sup><br>Complex: 48nM<sup>105</sup> | 17nM<sup>66</sup> | 93nM<sup>66</sup> |
| Important cargo (according to SGD yeastgenome.org) | Proteins, RNAs, and ribosomal subunits | Nuclear proteins, ribosome-bound peptides | Nuclear proteins, ribosome-bound peptides | Nab2, Hrp1, RNA export splicing machinery | Pdr1p, Yap1p, Ste12p, and Aft1p |

**Table S2.** Steady state localization of transport reporters for nuclear transport by Crm1, Kap95, Kap104, and Pse1in presence of PR_50_ or GA_50_

|  |  |  |  |  |  |
| --- | --- | --- | --- | --- | --- |
|  | Wild type | PR<sub>50</sub> |  | GA<sub>50</sub> |  |
| Transport reporters | N/C<sup>*1</sup><br>(no of cells) | N/C<br>(no of cells) | Fold change to WT (log2)<br>(p-value)<sup>*2</sup> | N/C<br>(no of cells) | Fold change to WT (log2)<br>(p-value)<sup>*2</sup> |
| No NLS | 1.24 ± 0.14<br>(178) | 1.24 ± 0.15<br>(228) | 0.01<br>(ns) | 1.11 ± 0.11<br>(115) | -0.16<br>($<0.0001$) |
| Ssb1 NES (Crm1) | 0.53 ± 0.15<br>(191) | 0.47 ± 0.16<br>(157) | -0.16<br>(0.0025) | 0.62 ± 0.22<br>(133) | 0.22<br>($<0.0001$) |
| Sv40 NLS (Kap60/Kap95) | 3.21 ± 0.89<br>(156) | 4.11 ± 1.44<br>(207) | 0.36<br>($<0.0001$) | 1.66 ± 0.99<br>(108) | -0.96<br>($<0.0001$) |
| Npm1 NLS (Kap60/Kap95) | 6.93 ± 3.37<br>(146) | 6.07 ± 4.99<br>(130) | -0.19<br>(ns) | 6.91 ± 5.52<br>(142) | 0.01<br>(ns) |
| Nab2 NLS (Kap104) | 7.59 ± 2.82<br>(105) | 6.37 ± 2.42<br>(112) | -0.25<br>($<0.0001$) | 5.25 ± 1.91<br>(105) | -0.51<br>($<0.0001$) |
| Pho4 NLS (Pse1) | 5.37 ± 2.71<br>(158) | 2.82 ± 1.10<br>(103) | -0.93<br>($<0.0001$) | 5.28 ± 2.19<br>(151) | -0.01<br>(ns) |
| <sup>*1</sup> N/C: fluorescence in nucleus over fluorescence in cytosol; median ± standard deviation | | | | | |
| <sup>*2</sup> ns= nonsignificant. Mann-Whitney test |  |  |  |  |  |

**Table S3.** Efflux rate constants of transport reporters in presence of PR_50_ or GA_50_

|  |  |  |  |  |  |
| --- | --- | --- | --- | --- | --- |
|  | Wild type | PR<sub>50</sub> |  | GA<sub>50</sub> |  |
| Transport reporters | Efflux rate constant<sup>*1</sup><br>(no of cells) | Efflux rate constant<sup>*1</sup><br>(no of cells) | Fold change to WT<br>(p-value)<sup>*2</sup> | Efflux rate constant<sup>*1</sup><br>(no of cells) | Fold change to WT (log2)<br>(p-value)<sup>*2</sup> |
| Sv40 NLS<br>(Kap60/Kap95) | 0.003 ± 0.001<br>(126) | 0.002 ± 0.003<br>(125) | (<0.0001) | ND | ND |
| Nab2 NLS<br>(Kap104) | 0.001 ± 0.003<br>(254) | 0.000 ± 0.001<br>(125) | (<0.0001) | 0.004 ± 0.005<br>(139) | (<0.0001) |
| Pho4 NLS<br>(Pse1) | 0.003 ± 0.004<br>(188) | 0.002 ± 0.003<br>(224) | (<0.0001) | 0.005 ± 0.004<br>(124) | (0.0001) |
| *<sup>1</sup> Median ± standard deviation *<sup>2</sup> ns= nonsignificant, Mann-Whitney test *<sup>3</sup> ND: not determined |  |  |  |  |  |

**Table S4.**
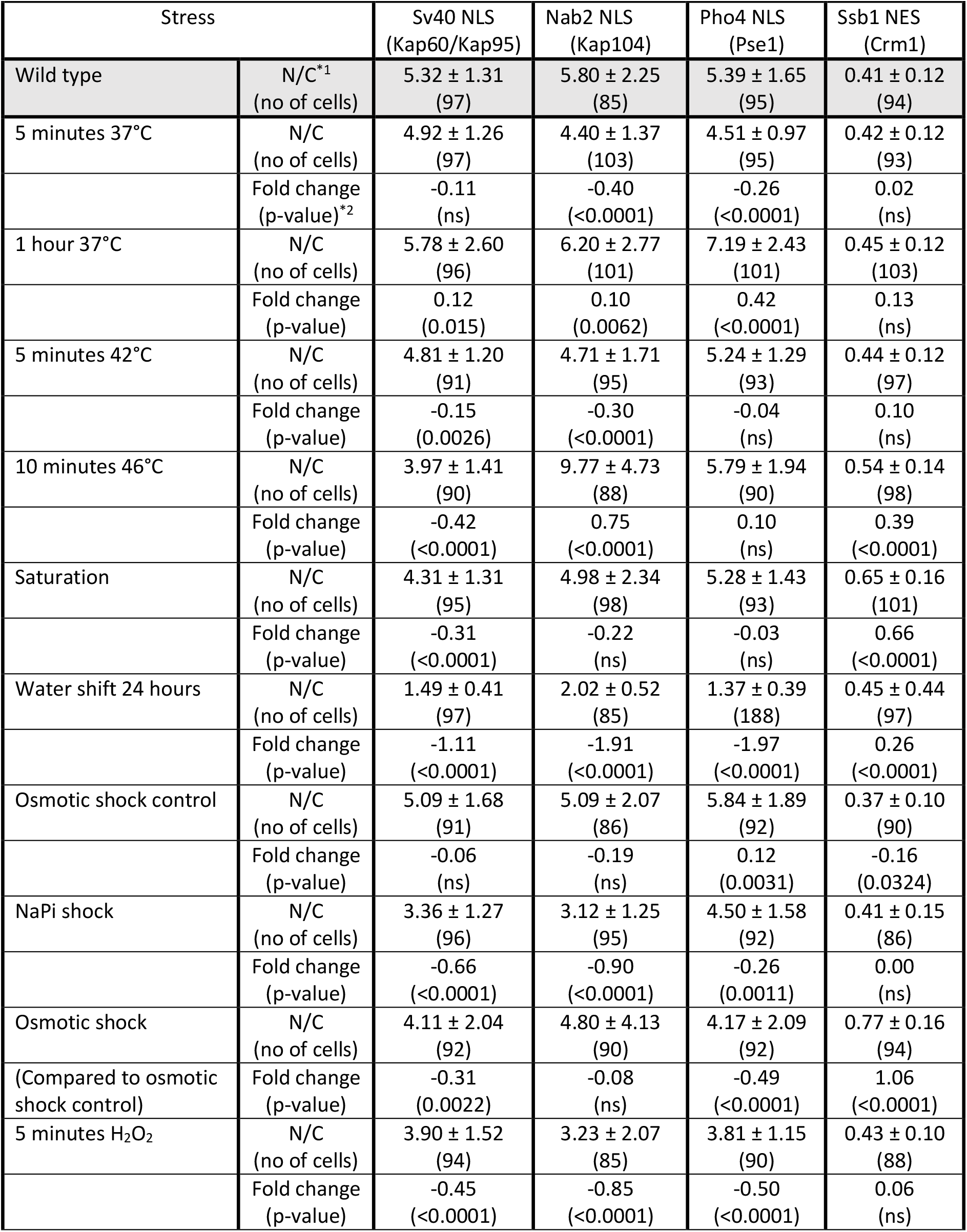
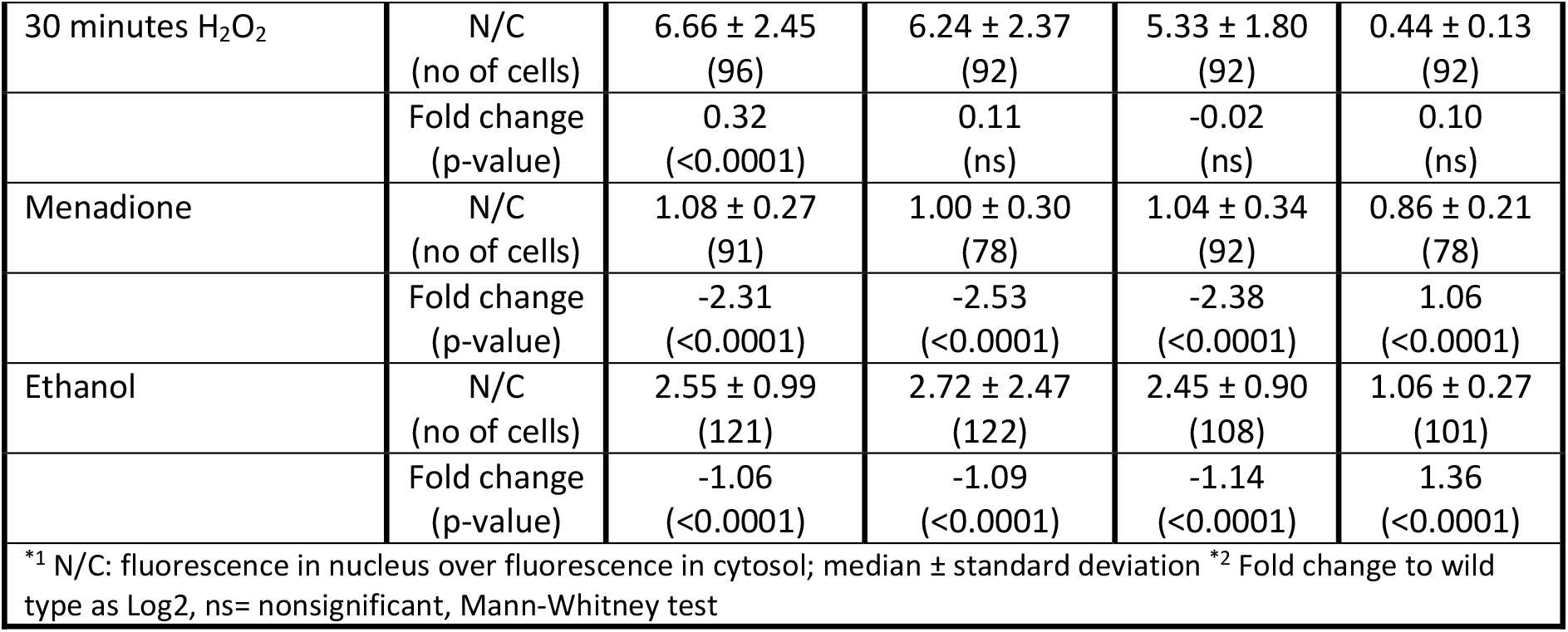
Steady state localization of transport reporters for nuclear transport by Crm1, Kap95, Kap104, and Pse1 under stress conditions

| Stress |  | Sv40 NLS<br>(Kap60/Kap95) | Nab2 NLS<br>(Kap104) | Pho4 NLS<br>(Pse1) | Ssb1 NES<br>(Crm1) |
| --- | --- | --- | --- | --- | --- |
| Wild type | N/C*<sup>1</sup><br>(no of cells) | 5.32 ± 1.31<br>(97) | 5.80 ± 2.25<br>(85) | 5.39 ± 1.65<br>(95) | 0.41 ± 0.12<br>(94) |
| 5 minutes 37°C | N/C<br>(no of cells) | 4.92 ± 1.26<br>(97) | 4.40 ± 1.37<br>(103) | 4.51 ± 0.97<br>(95) | 0.42 ± 0.12<br>(93) |
|  | Fold change<br>(p-value)*<sup>2</sup> | -0.11<br>(ns) | -0.40<br>(<0.0001) | -0.26<br>(<0.0001) | 0.02<br>(ns) |
| 1 hour 37°C | N/C<br>(no of cells) | 5.78 ± 2.60<br>(96) | 6.20 ± 2.77<br>(101) | 7.19 ± 2.43<br>(101) | 0.45 ± 0.12<br>(103) |
|  | Fold change<br>(p-value) | 0.12<br>(0.015) | 0.10<br>(0.0062) | 0.42<br>(<0.0001) | 0.13<br>(ns) |
| 5 minutes 42°C | N/C<br>(no of cells) | 4.81 ± 1.20<br>(91) | 4.71 ± 1.71<br>(95) | 5.24 ± 1.29<br>(93) | 0.44 ± 0.12<br>(97) |
|  | Fold change<br>(p-value) | -0.15<br>(0.0026) | -0.30<br>(<0.0001) | -0.04<br>(ns) | 0.10<br>(ns) |
| 10 minutes 46°C | N/C<br>(no of cells) | 3.97 ± 1.41<br>(90) | 9.77 ± 4.73<br>(88) | 5.79 ± 1.94<br>(90) | 0.54 ± 0.14<br>(98) |
|  | Fold change<br>(p-value) | -0.42<br>(<0.0001) | 0.75<br>(<0.0001) | 0.10<br>(ns) | 0.39<br>(<0.0001) |
| Saturation | N/C<br>(no of cells) | 4.31 ± 1.31<br>(95) | 4.98 ± 2.34<br>(98) | 5.28 ± 1.43<br>(93) | 0.65 ± 0.16<br>(101) |
|  | Fold change<br>(p-value) | -0.31<br>(<0.0001) | -0.22<br>(ns) | -0.03<br>(ns) | 0.66<br>(<0.0001) |
| Water shift 24 hours | N/C<br>(no of cells) | 1.49 ± 0.41<br>(97) | 2.02 ± 0.52<br>(85) | 1.37 ± 0.39<br>(188) | 0.45 ± 0.44<br>(97) |
|  | Fold change<br>(p-value) | -1.11<br>(<0.0001) | -1.91<br>(<0.0001) | -1.97<br>(<0.0001) | 0.26<br>(<0.0001) |
| Osmotic shock control | N/C<br>(no of cells) | 5.09 ± 1.68<br>(91) | 5.09 ± 2.07<br>(86) | 5.84 ± 1.89<br>(92) | 0.37 ± 0.10<br>(90) |
|  | Fold change<br>(p-value) | -0.06<br>(ns) | -0.19<br>(ns) | 0.12<br>(0.0031) | -0.16<br>(0.0324) |
| NaPi shock | N/C<br>(no of cells) | 3.36 ± 1.27<br>(96) | 3.12 ± 1.25<br>(95) | 4.50 ± 1.58<br>(92) | 0.41 ± 0.15<br>(86) |
|  | Fold change<br>(p-value) | -0.66<br>(<0.0001) | -0.90<br>(<0.0001) | -0.26<br>(0.0011) | 0.00<br>(ns) |
| Osmotic shock | N/C<br>(no of cells) | 4.11 ± 2.04<br>(92) | 4.80 ± 4.13<br>(90) | 4.17 ± 2.09<br>(92) | 0.77 ± 0.16<br>(94) |
| (Compared to osmotic shock control) | Fold change<br>(p-value) | -0.31<br>(0.0022) | -0.08<br>(ns) | -0.49<br>(<0.0001) | 1.06<br>(<0.0001) |
| 5 minutes H<sub>2</sub>O<sub>2</sub> | N/C<br>(no of cells) | 3.90 ± 1.52<br>(94) | 3.23 ± 2.07<br>(85) | 3.81 ± 1.15<br>(90) | 0.43 ± 0.10<br>(88) |
|  | Fold change<br>(p-value) | -0.45<br>(<0.0001) | -0.85<br>(<0.0001) | -0.50<br>(<0.0001) | 0.06<br>(ns) |
| 30 minutes H<sub>2</sub>O<sub>2</sub> | N/C<br>(no of cells) | 6.66 ± 2.45<br>(96) | 6.24 ± 2.37<br>(92) | 5.33 ± 1.80<br>(92) | 0.44 ± 0.13<br>(92) |
| | Fold change<br>(p-value) | 0.32<br>($<0.0001$) | 0.11<br>(ns) | -0.02<br>(ns) | 0.10<br>(ns) |
| Menadione | N/C<br>(no of cells) | 1.08 ± 0.27<br>(91) | 1.00 ± 0.30<br>(78) | 1.04 ± 0.34<br>(92) | 0.86 ± 0.21<br>(78) |
| | Fold change<br>(p-value) | -2.31<br>($<0.0001$) | -2.53<br>($<0.0001$) | -2.38<br>($<0.0001$) | 1.06<br>($<0.0001$) |
| Ethanol | N/C<br>(no of cells) | 2.55 ± 0.99<br>(121) | 2.72 ± 2.47<br>(122) | 2.45 ± 0.90<br>(108) | 1.06 ± 0.27<br>(101) |
| | Fold change<br>(p-value) | -1.06<br>($<0.0001$) | -1.09<br>($<0.0001$) | -1.14<br>($<0.0001$) | 1.36<br>($<0.0001$) |
| *<sup>1</sup> N/C: fluorescence in nucleus over fluorescence in cytosol; median ± standard deviation *<sup>2</sup> Fold change to wild type as Log2, ns= nonsignificant, Mann-Whitney test |  |  |  |  |  |

**Table S5.** Steady state localization of transport reporters for nuclear transport by Crm1, Kap95, Kap104, and Pse1 cargos in mutants

|  |  |  |  |  |  |
| --- | --- | --- | --- | --- | --- |
| Stress |  | Sv40 NLS<br>(Kap60/Kap95) | Nab2 NLS<br>(Kap104) | Pho4 NLS<br>(Pse1) | Ssb1 NES<br>(Crm1) |
| Sxm1Δ | N/C<br>(no of cells) | 5.82 ± 2.21<br>(102) | 6.56 ± 2.14<br>(100) | 4.36 ± 1.20<br>(102) | 0.40 ± 0.11<br>(94) |
| | Fold change<br>(p-value)*<sup>2</sup> | 0.39<br>($<0.0001$) | -0.21<br>(0.0257) | -0.31<br>($<0.0001$) | 0.16<br>(0.0065) |
| Kap114Δ | N/C<br>(no of cells) | 5.01 ± 1.82<br>(98) | 7.02 ± 4.44<br>(100) | 4.71 ± 2.20<br>(102) |  |
| | Fold change<br>(p-value) | 0.64<br>($<0.0001$) | -0.11<br>(ns) | -0.19<br>(ns) | |
| Kap120Δ | N/C<br>(no of cells) | 5.39 ± 2.00<br>(78) | 9.46 ± 3.98<br>(148) | 7.30 ± 2.91<br>(153) | 0.38 ± 0.12<br>(94) |
| | Fold change<br>(p-value) | 0.75<br>($<0.0001$) | 0.32<br>($<0.0001$) | 0.44<br>($<0.0001$) | 0.30<br>($<0.0001$) |
| Kap122Δ | N/C<br>(no of cells) | 4.88 ± 2.00<br>(125) | 4.21 ± 1.32<br>(209) | 3.48 ± 0.98<br>(135) | 0.39 ± 0.14<br>(96) |
| | Fold change<br>(p-value) | 0.61<br>($<0.0001$) | -0.70<br>(0.0015) | -0.20<br>(0.0048) | 0.30<br>($<0.0001$) |
| Kap123Δ | N/C<br>(no of cells) | 4.04 ± 1.10<br>(73) | 8.65 ± 2.26<br>(149) | 3.29 ± 0.75<br>(110) | 0.33 ± 0.07<br>(102) |
| | Fold change<br>(p-value) | 0.33<br>($<0.0001$) | -0.65<br>($<0.0001$) | -0.81<br>($<0.0001$) | 0.13<br>(0.0144) |
| RanGAP<br>overexpression | N/C<br>(no of cells) | 4.35 ± 2.05<br>(100) | 2.86 ± 1.59<br>(98) |  | 0.46 ± 0.13<br>(91) |
| | Fold change<br>(p-value) | -0.29<br>(0.0008) | -1.02<br>($<0.0001$) | | 0.16<br>(ns) |
| RanGEF<br>overexpression | N/C<br>(no of cells) | 4.00 ± 2.02<br>(97) | 5.15 ± 1.67<br>(91) | 4.35 ± 1.87<br>(95) | 0.46 ± 0.15<br>(88) |
| | Fold change<br>(p-value) | -0.41<br>($<0.0001$) | -0.17<br>(0.0257) | -0.31<br>($<0.0001$) | 0.16<br>(0.0065) |
| *<sup>1</sup> N/C: fluorescence in nucleus over fluorescence in cytosol; median ± standard deviation *<sup>2</sup> Fold change to wild type as Log2, ns= nonsignificant, Mann-Whitney test |  |  |  |  |  |

**Fig S2.**
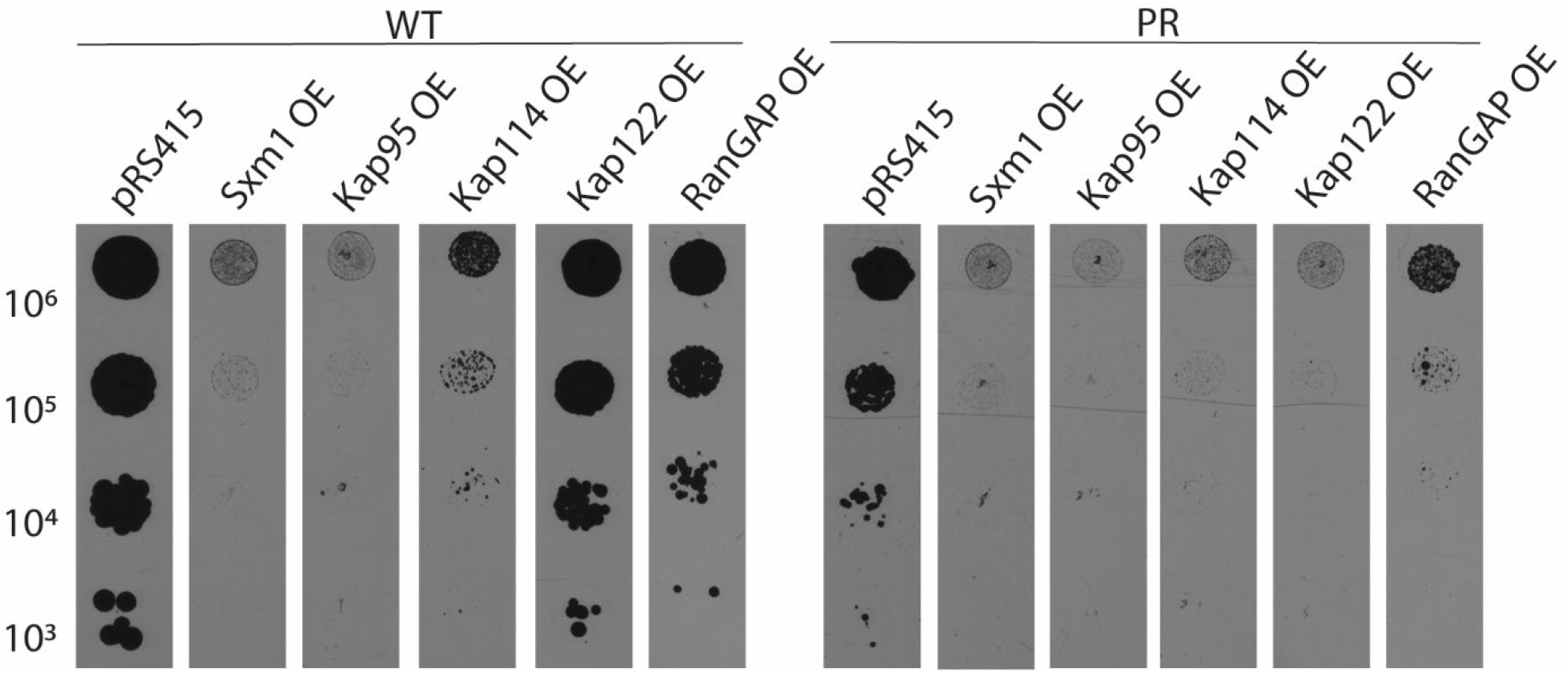
Overexpression of NTRs is toxic to cells, additionally so with co-expression of PR_50_. **A)** Galactose induced overexpression of NTRs Sxm1, Kap95, Kap114 is toxic to cells; overexpression of Kap122, and RanGAP, is less toxic (left panel). Simultaneous galactose induced co-expression of PR_50_, is lethal for all NTRs and RanGAP OE.

